# Endonuclease activity of RecJ from extremely alkaliphilic Bacillus alcalophilus

**DOI:** 10.1101/2020.02.19.956169

**Authors:** Minggang Zheng, Wen Wang, Liya Ma, Ling Wang, Lingyun Qu, Xipeng Liu, Hailiang Liu

## Abstract

At present, all documented RecJs are exonucleases, degrading single-stranded nucleic acids. Here, we report a novel RecJ, from the extremely alkaliphilic bacterium *Bacillus alcalophilus* (BaRecJ), which possesses endonuclease activity and can cleave supercoiled DNA. BaRecJ contains the typical DHH and DHHA1 domains, which are conserved in all RecJs, and a functionally unknown PIWI-like domain at the C-terminus. The endonuclease activity originates from the C-terminal domain of BaRecJ which contains PIWI-like domain, and the exonuclease activity from the DHH and DHHA1 domains. Mutational analysis reveals that several important residues affect the endonuclease activity of BaRecJ. Moreover, BaRecJ cleaves specific target sequences at moderate temperature when directed by a phosphorothioate-modified single-stranded DNA (S-modified ssDNA) guide. These findings suggest that BaRecJ is substantially different from any reported RecJs and has the potential to be developed as a new gene editing tool.

---

As a DHH phosphatase superfamily member, RecJ has long been considered to function on ssDNA or ssRNA as an exonuclease (1, 2), and is involved in diverse processes including resecting DNA ends in homologous recombination (HR), degrading the 5ʹ- strand of the DNA unwound by RecQ helicase at double-stranded DNA breaks (3) mediating the excision step (4), reducing homology-facilitated illegitimate recombination events (5), and rescuing stalled replication forks (6, 7). RecJ-like can also degrade ssRNA from the 3ʹ-end, and proofread 3ʹ-mismatched RNA primers in DNA replication (8).

The enzymatic functions and activities of proteins are greatly influenced by the domains that they contain. RecJ from the bacterium *Escherichia coli* (EcRecJ), a typical RecJ, possesses DHH, DHHA1 and OB-fold domains, and exhibits 5ʹ- to 3ʹ- ssDNA exonuclease activity (9). *Thermus thermophilus* RecJ (TtRecJ) and *Deinococcus radiodurans* RecJ (DrRecJ) possess the aforementioned domains and an additional C-terminal element, which contributes to their DNA binding capability (10,11,12). RecJ from the archaeon *Pyrococcus furiosus* can degrade ssRNA from both the 5ʹ- and 3ʹ- ends, despite only possessing DHH and DHHA1 domains^8^. In contrast, Cdc45, a eukaryotic RecJ homolog, contains DHH and DHHA domains and an additional C-terminal domain; it lacks nuclease activity but has ssDNA binding capability (13). Although RecJ homologous genes have been found in almost all bacteria, archaea and eukaryotes, RecJs have only been shown to degrade ssDNA and ssRNA as exonucleases.

The crystal structures of RecJs from *T. thermophilus* and *D. radiodurans* have been solved. The structure of TtRecJ reveals a pocket, formed from the OB fold domain. The hole near the active site is too narrow for double-stranded (ds) DNA to fit, ensuring the ssDNA-specificity of TtRecJ (7, 14). The structure of DrRecJ shows that the terminus of ssDNA is anchored to the 5ʹ- phosphate binding pocket above the active site, which is guarded by a helical gateway. This prevents dsDNA from entering the active site, and the OB fold domain is located beside the DNA entrance. The structure suggests that a free 5ʹ- ssDNA is essential for DrRecJ nuclease activity (15).

In this study, we first report endonuclease activity of *Bacillus alcalophilus* RecJ (BaRecJ; WP_003324372.1), and characterize its guide-dependent and guide-independent cleavage reactions. Obviously different from reported RecJs, other than in the conserved DHH and DHHA1 domains, this protein contains a functionally unknown PIWI-like domain (residues 471–653) at the C-terminus; such a domain is widely present in RecJs of members of the genus *Bacillus*. *In vivo*, the expression of BaRecJ affects host cell morphology, growth and survival, and causes a decrease of genomic and plasmid DNA content. Based on this finding, the enzymatic properties of BaRecJ were biochemically characterized *in vitro*. The results show that BaRecJ cleaves supercoiled plasmid DNA into open circular or linear DNA, or even completely degrades it into nucleotides, indicating its function as a DNA endonuclease and exonuclease. Further analysis of nuclease activities confirmed that the endonuclease activity originates from the C-terminal domain of BaRecJ which contains PIWI-like domain, and the exonuclease activity from the DHH and DHHA1 domains. We also found that S-modified ssDNA can be a guide in guide-dependent DNA cleavage by BaRecJ. Taken together, our findings suggest that the newly identified BaRecJ is substantially different from other reported RecJ proteins. On one hand, it is probably involved in homologous recombination and mismatch repair based on exonuclease activity, and on the other hand, it functions as an endonuclease like Argonaute (Ago) proteins to perform silencing of foreign genes.

## RESULTS AND DISCUSSION

### Expression of BaRecJ decreases transformation efficiency

As a multidomain protein, RecJ has conserved DHH and DHHA1 domains, which are the catalytic core for RecJ exonuclease activity (16) (Figure 1a and Supplementary Figure 1). Some RecJs also possess a C-terminal domain, and these C-terminal domains confer diverse functions (7, 11). In addition to the DHH and DHHA1 domains, BaRecJ possesses a PIWI-like domain at its C-terminus (Figure 1a). The PIWI-like domain possesses endonuclease activity and plays an essential role in RNA-induced silencing (17, 18).

**Figure 1.**
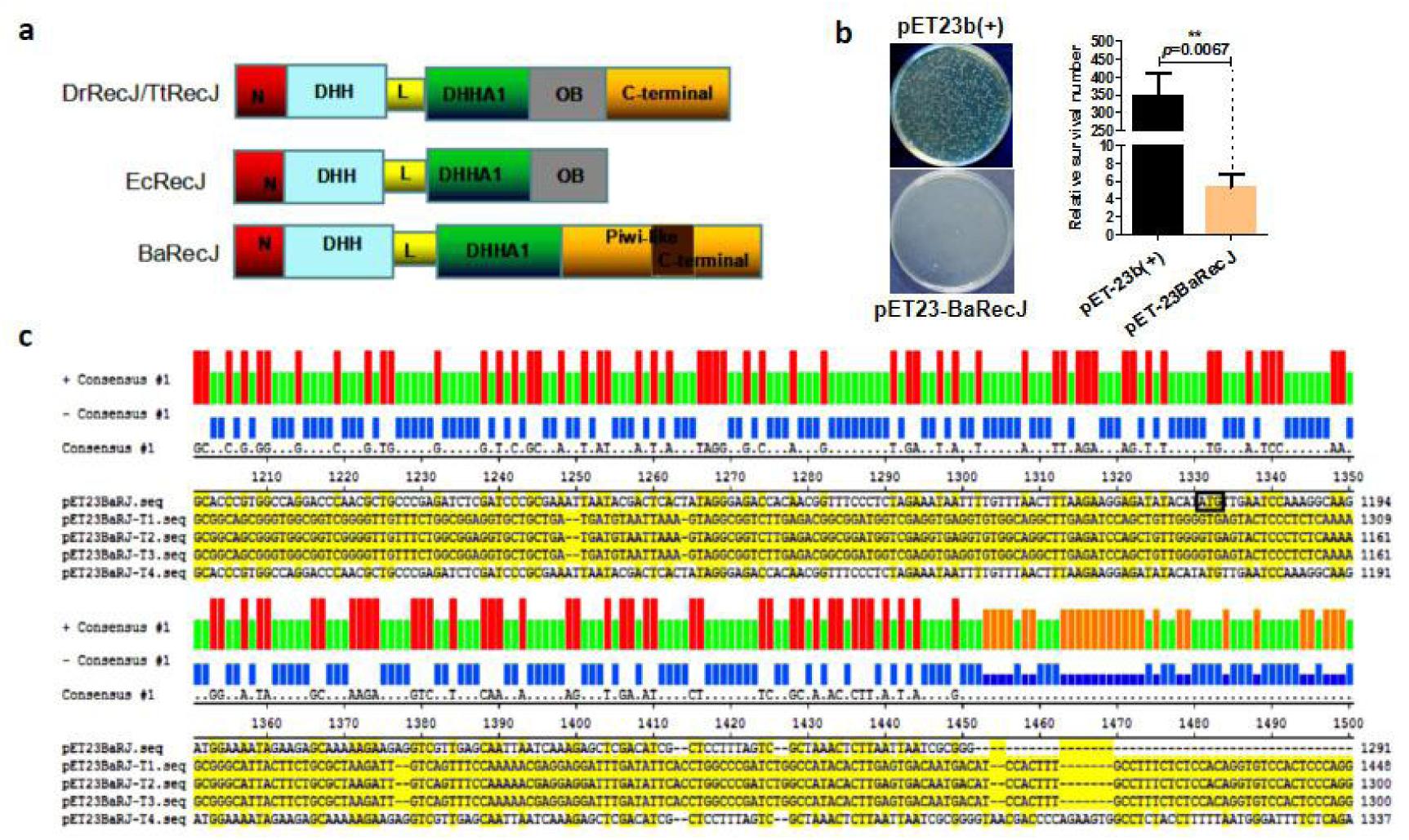
Effect of *Bacillus alcalophilus* RecJ (BaRecJ) expression on plasmid DNA in *Escherichia coli*. a Schematic representation of RecJ family members. Dr, Tt, Ec and Ba are *Deinococcus radiodurans*, *Thermus thermophilus*, *E. coli* and *B. alcalophilus*, respectively. N, DHH, DHHA1, OB, PIWI-like and C-terminal denote the main structural domains; L is a linker. The dark shaded area shows the sequence overlap between the PIWI-like domain and the C-terminal domain in BaRecJ. b *E. coli* BL21 (DE3) harboring pET23b(+) and pET23BaRJ were respectively cultured overnight. Appropriate volumes of cultures were spread onto plates containing 50 μg ml^−1^ ampicillin and incubated at 37°C overnight. The Figure shows comparison of the transformation efficiency of *E. coli* BL21 (DE3) transformed with either pET-23b(+) or pET23BaRecJ (Supplementary Data Table 1). Error bars show standard deviations. c Sequence alignment of pET23BaRecJ with sequences of pET23BaRecJT1 to T4; the plasmids pET23BaRecJT1 to T4 were respectively isolated from transformants surviving on the ampicillin-containing agar plates. ATG, the start codon of the *BarecJ* gene, is boxed. Red columns indicate unmutated bases (identical between the five plasmids), orange and green columns denote mutated bases.

To elucidate the functions of RecJ from *B. alcalophilus*, we cloned and expressed its gene. We found that the transformation efficiency in *E. coli* BL21 (DE3) was very low when using a constitutive expression vector carrying *BarecJ* (around 5.3% compared with empty vector pET-23b (+); *P* < 0.0067) (Figure 1b). What caused the low transformation efficiency? We found that plasmids isolated from the above transformants showed profound variation, except in a few regions essential for copy number control and antibiotic resistance (Figure 1c, Supplementary Figure 2 and Supplementary Table 1). Previously reported RecJs only have exonuclease activity, which cannot lead to decreased transformation efficiency (19–22). So, we suggest that BaRecJ has a function different from that of any reported RecJs. It performs endonuclease activity and cleaves at least the plasmid DNA to decrease the tranformation efficiency.

### Expression of BaRecJ decreases host cell survival rate and DNA content

Induced expression of BaRecJ in *E. coli* BL21 (DE3) was carried out to verify its effect on host cells. Isopropyl-β-D-thiogalactoside (IPTG) induction of BaRecJ expression caused the cells to form long filaments (Figure 2a, upper right panel), similar to cells during inhibition of cell division (23). In contrast, induction of expression did not lead to cells harboring control plasmid showing abnormal cell morphology (Figure 2a, lower right panel). The survival rate of cells transformed with the BaRecJ expression vector was 1.3% of that of cells containing empty vector on plates containing 0.1 mM IPTG (*P* < 0.0001; Figure 2b). In liquid medium, it was found that the surviving number of cells containing BaRecJ decreased by 99% in 4.0 h, while it increased 108-fold for cells containing empty vector (Figure 2c). In addition, we found that the plasmid and genomic DNA content in cells carrying BaRecJ was lower than that in cells transformed with control vector after induction for 3.0 h (*P* = 0.0092, Figure 2d, and *P* = 0.0076, Figure 2e). Similar results were obtained with BhRecJ (the RecJ homologue from *B. halodurans*), BoRecJ (from *B. okhensis*), BcRecJ (from *B. cereus*), and FeRecJ (from *Fictibacillus enclensis*) (Supplementary Figure 3). These results imply that BaRecJ may cause double strand breaks (DSBs) in plasmid and genomic DNA in *E. coli* host cells, and lead to a decrease in cellular DNA content and cell death.

**Figure 2.**
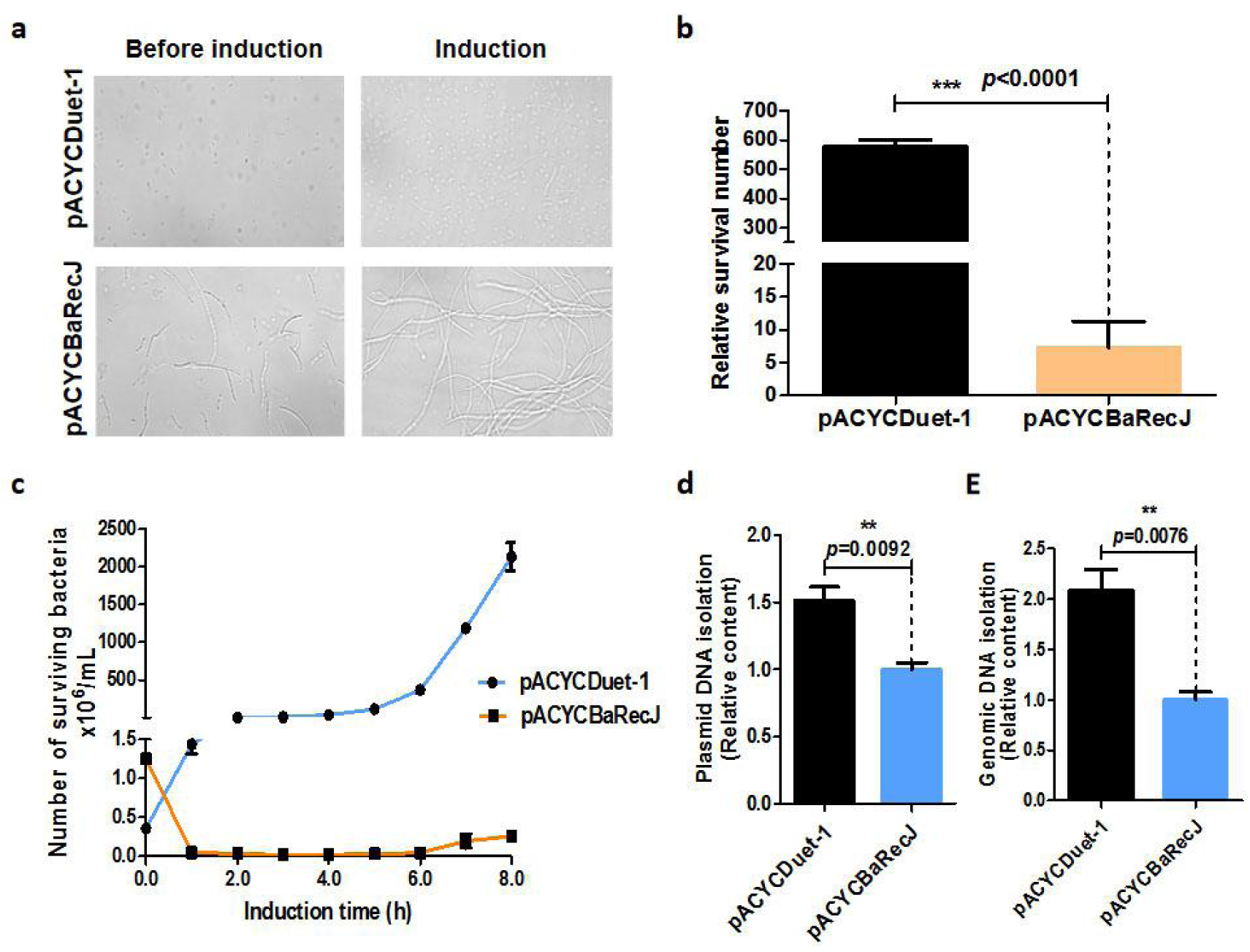
Expression of BaRecJ decreases *E. coli* cell survival rate and changes cell morphology. a Differences in morphology of *E. coli* BL21 (DE3) transformants with pACYCDuet-1 (lower) and pACYCBaRecJ (upper). The cultures were imaged under a microscope (40×) without (left panel) isopropyl-β-D-thiogalactoside (IPTG) induction or with (right panel) IPTG induction, respectively. b Comparison of surviving numbers of transformants with pACYCBaRecJ or pACYCDuet-1. All transformants were cultivated overnight with IPTG induction. c Survival curves of transformants with pACYCBaRecJ and pACYCDuet-1, respectively. d Plasmid and genomic DNAs were extracted from the same amount of biomass of *E. coli* induced for 3 h, respectively. Error bars depict standard deviations of biological triplicates.

### Characterization of the nuclease activity of BaRecJ

To confirm that the effect of BaRecJ toward host cells is due to nuclease activity, we purified BaRecJ protein after inducing expression in *E. coli* at 16°C (Figure 3 a, d). At this temperature, the nuclease activity of BaRecJ was inhibited. *In vitro*, it was found that BaRecJ can degrade supercoiled plasmid DNAs into open circular or linear ones at 23 to 60°C (Figure 3d), and the amount of BaRecJ had a tremendous impact on the degradation efficiency (Figure 3b). When the digestion time or the amount of BaRecJ was increased, a smear could be generated on electrophoresis of BaRecJ-digested plasmid DNA. Supercoiled plasmid was almost completely digested within 2 h when the concentration ratio of enzyme to plasmid substrate was 20:1 (Figure 3c). Digestion was promoted by both Mg^2+^ and Mn^2+^ (Figure 3e and Supplementary Figure 4), and not sensitive to Cu^2+^ (Supplementary Figure 4), which is similar to findings for many documented RecJ nucleases. But different with other RecJs, the digestion of BaRecJ is not sensitive to Co^2+^ (14, 24). BaRecJ can digest many vectors with different sequences at different rates, including pGADT7, pET-28, pUC57, pTARGET and pCRCT (Figure 3f). BaRecJ can also digest linear dsDNA and generate a smear in electrophoresis (Supplementary Figure 5).

**Figure 3.**
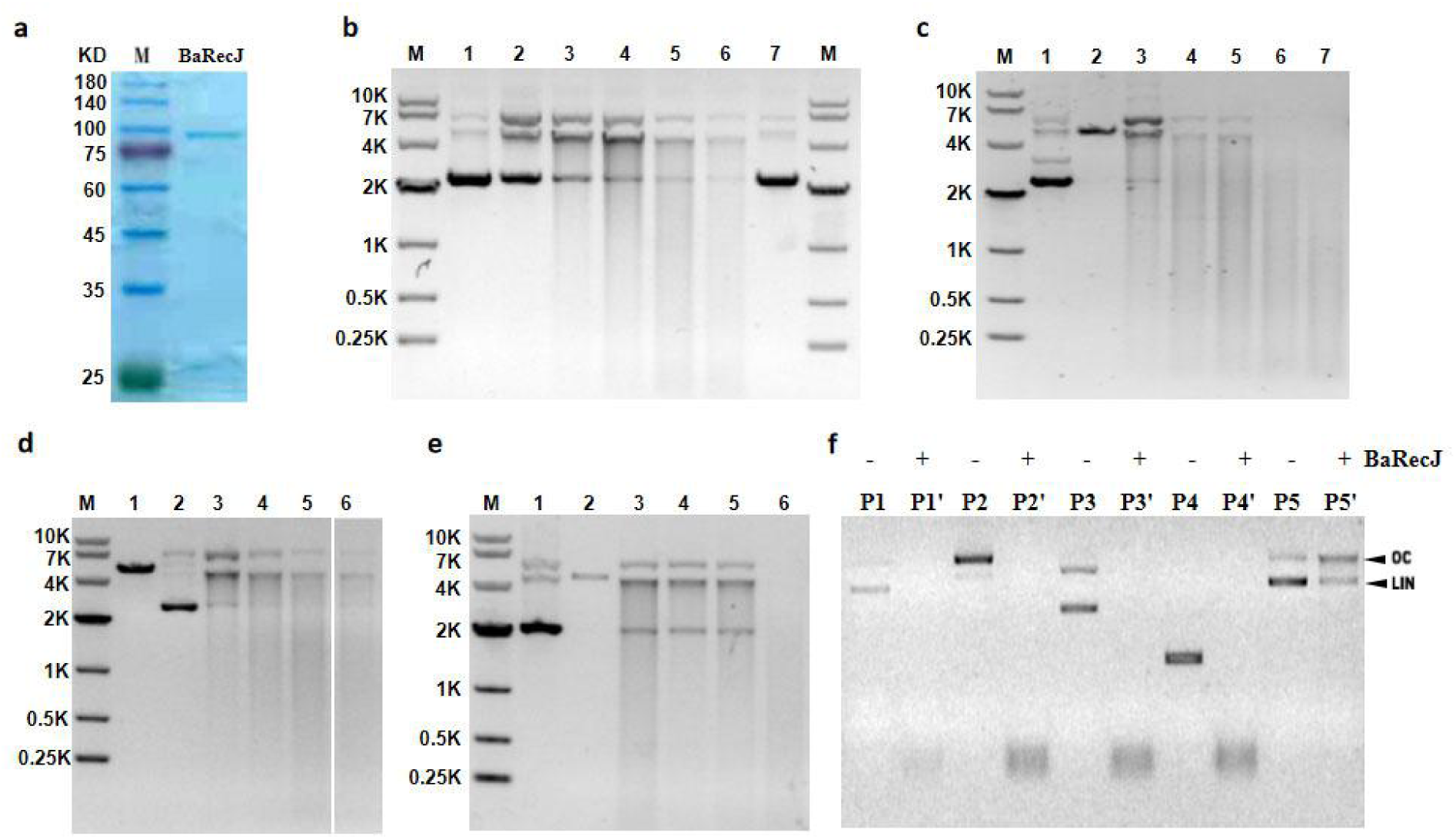
Characterization of the nuclease activity of BaRecJ toward plasmid DNA. **a** Purity analysis of recombinant RecJs by 15% SDS-PAGE. **b** Titration of BaRecJ with pUC19-s. BaRecJ was incubated with vector pUC19-s at 37°C for 2 h. The molar ratios of BaRecJ to pUC19-s were 1:1, 5:1, 10:1, 20:1, and 40:1 with enzyme concentration 250 nM (lanes 2–6). **c** Time course of BaRecJ activity toward pUC19-s at 37°C. The molar ratio of BaRecJ to pUC19-s was 20:1 with enzyme concentration 500 nM. Lane 1: pUC19-s control linearized with *Eco*RI. Lane 2: pUC19-s control plasmid. Lanes 3–7: pUC19-s digested for 30, 60, 90, 120, and 180 min by BaRecJ, respectively. **d** Effect of temperature on the activity of BaRecJ toward pUC19-s. The molar ratio of BaRecJ to pUC19-s was 20:1 with enzyme concentration 500 nM. The incubation time was 2 h. Lanes 3–6: 23, 30, 37 and 60°C, respectively. **e** Effect of Mg^2+^ on the activity of BaRecJ. The reactions were performed at 37°C for 2 h with 0.1, 1, 5 and 10 mM Mg^2+^ (lanes 3–6, respectively). The molar ratio of BaRecJ to pUC19-s was 20:1 and the enzyme concentration was 500 nM. **f** Nuclease activity of BaRecJ toward various plasmids. The reactions were incubated at 37°C for 3 h. P1, P2, P3, P4, and P5 denote pGADT7, pET-28, pUC57, pTARGET, and pCRCT, respectively.

BaRecJ can digest the classical substrate ssDNA from the 5ʹ-terminus, and conserved residues are required for the ssDNA-specific 5ʹ-exonuclease activity. Like in classical RecJ (15), H170 is a key residue that was essential for this BaRecJ exonuclease activity (Figure 4a, left two panels, and Supplementary Table 4). The results of electrophoretic mobility shift assays (EMSAs) support the nuclease activity toward ssDNA substrates (Figure 4b). Among the experimental substrates, ssDNA was bound most strongly by BaRecJ (Figure 4b, left panel), which is consistent with the enzyme having its highest nuclease activity toward this substrate. It was further found that BaRecJ can bind dsDNA; however, it could not digest this substrate (Figure 4a, right two panels). These results imply that BaRecJ has some characteristics of both RecJ and Ago proteins; it has exonuclease activity like a typical RecJ, and endonuclease activity like a prokaryotic Ago (pAgo) protein.

**Figure 4.**
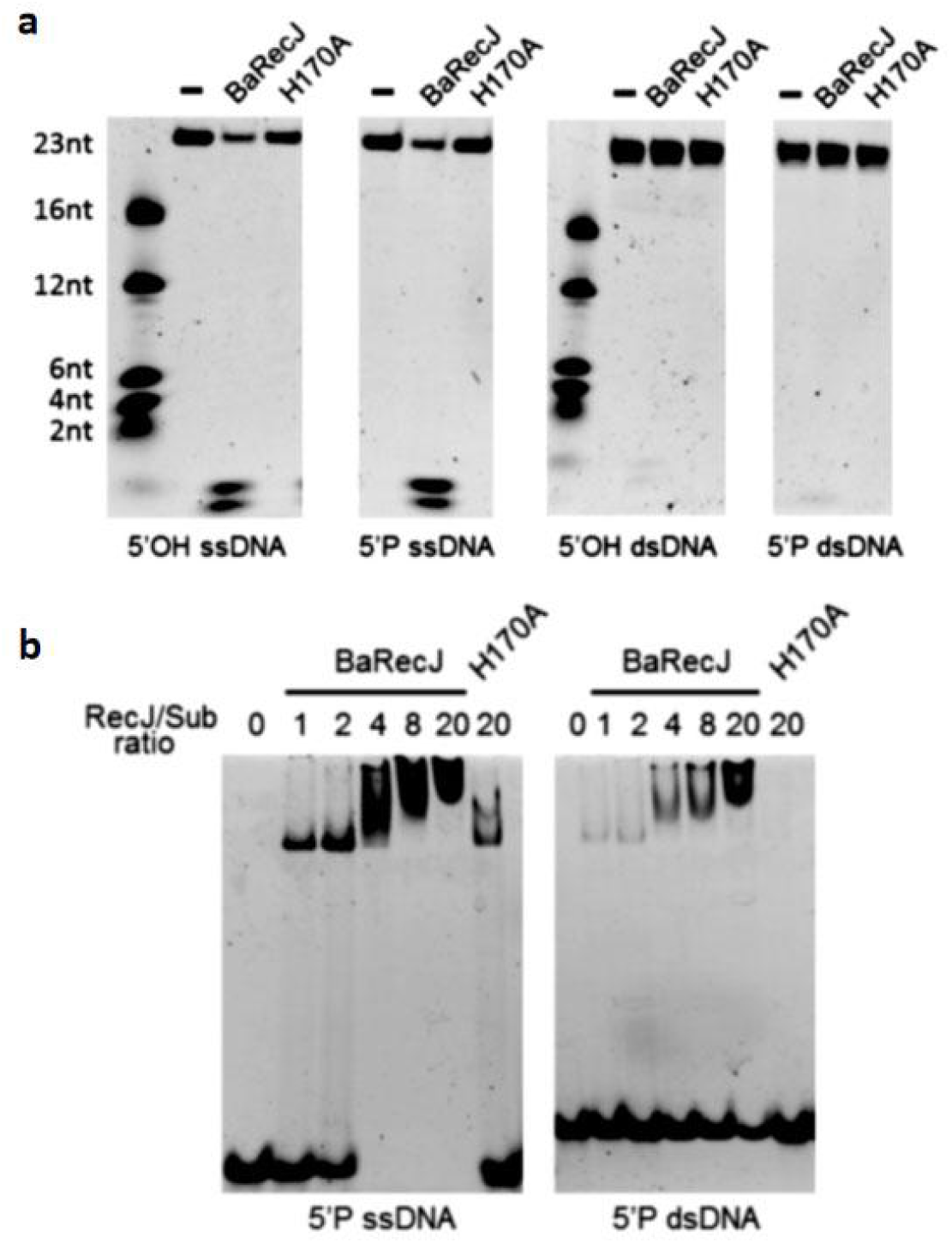
The exonuclease activity of BaRecJs toward single-stranded and double-stranded oligodeoxyribonucleotides. **a** Various oligodeoxyribonucleotides (2.5 pmol) in single-stranded or double-stranded form were incubated with 2.5 pmol enzyme at 37°C for 20 min. **b** Binding capability of BaRecJs to various oligodeoxyribonucleotides in single-stranded or double-stranded form. In electrophoretic mobility shift assays, Ca^2+^ was used in the reaction buffer instead of Mn^2+^ to prevent the cleavage of substrates.

### Mutational analysis of endonuclease activity

To determine the active site of BaRecJ endonuclease activity, we first used online prediction tools. BaRecJ spans residues 1–787, including DHH superfamily (1–550) and ssDNA exonuclease activity superfamily (550–787) domains, and the PIWI-like region near the C-terminus (471–653) (Figure 5a). We recombinantly expressed each of these fragments of the protein. *In vitro*, the fragment consisting of residues 471–787 retained endonuclease activity toward pUC19-s and pmg36e. It was further found that although the PIWI-like region had some activity toward pUC19-s and pmg36e, it was similar to that of the 653–787 amino acid (aa) segment, and weaker than that of the 471–787 aa segment (Figure 5b, c; Supplementary Figure 6). These data suggest that the DNA chopping activity does not depend on full-length BaRecJ, and the active site for this reaction is contained in the C-terminal fragment of the protein.

**Figure 5.**
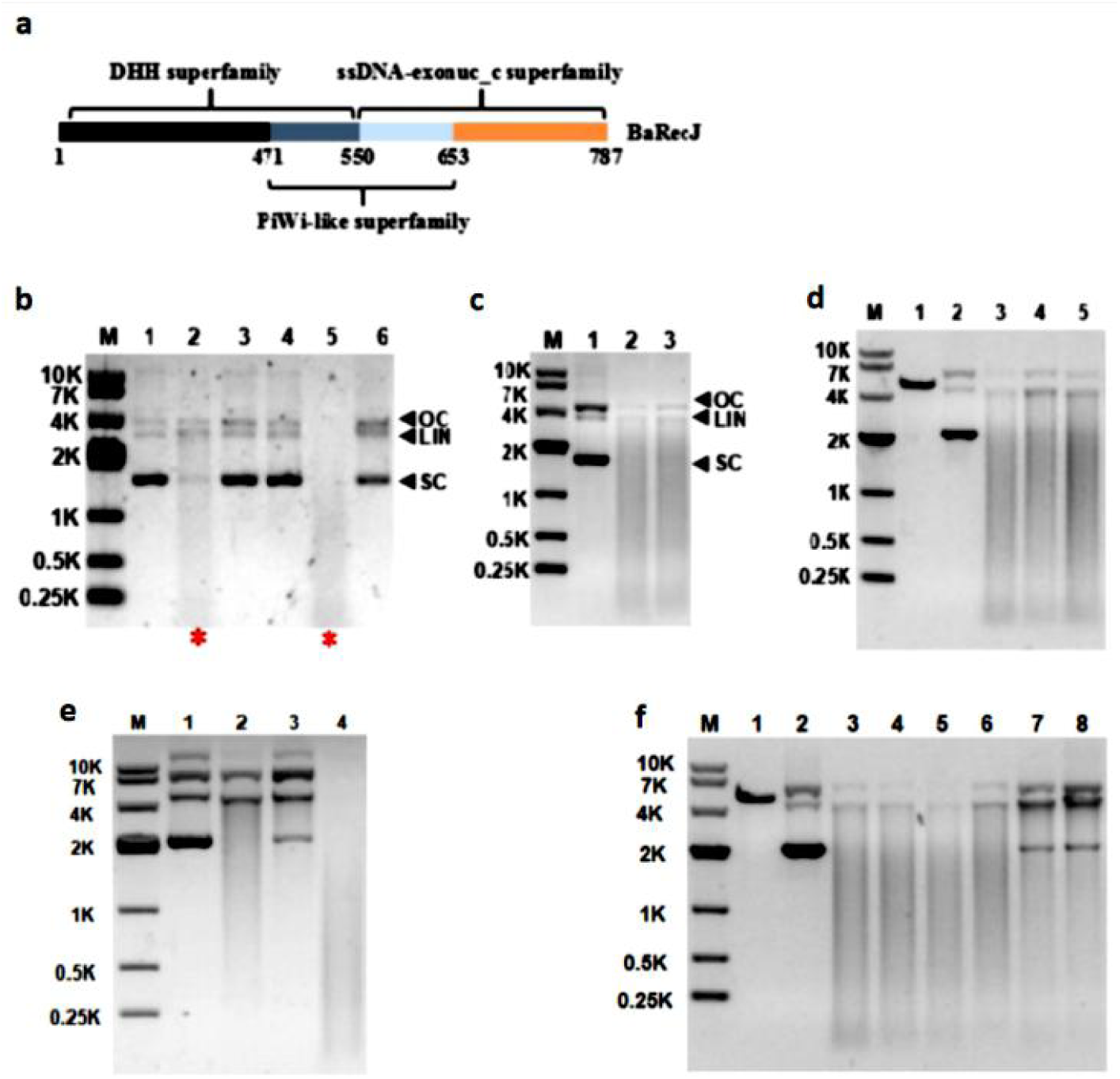
The endonuclease activity of truncations and point mutants of BaRecJ. **a** Schematic representation of BaRecJ. **b, c** Electrophoresis of the final plasmid degradation products of wild-type BaRecJ and its truncations (500 nM protein, 750 ng plasmid DNA, cleavage for 2 h at 37°C). (b), Lanes 1–6: pmg36e control plasmid, pmg36e cleaved by wt BaRecJ, and pmg36e cleaved by fragments of BaRecJ consisting of amino acids (aa) 1–241, aa 241–471, aa 471–787, and aa 1–471, respectively. (c), Lanes 1–3: pmg36e control plasmid, pmg36e cleaved by wt BaRecJ, and pmg36e cleaved by a fragment of BaRecJ consisting of aa 471–653. **d, e, f** Electrophoresis of the final plasmid degradation products of wt BaRecJ and its mutants (500 nM protein, 750 ng plasmid DNA, cleavage for 2 h at 37°C). (d), Lanes 1–5: pUC19-s linearized with *Eco*RI, pUC19-s control plasmid, pUC19-s cleaved by wt BaRecJ, and pUC19-s cleaved by BaRecJ E600A mutant or S762A mutant. (e), Lanes 1–4: pUC19-s control plasmid, pUC19-s cleaved by wt BaRecJ, and pUC19-s cleaved by BaRecJ D561A mutant or L634A mutant. (f), Lanes 1–8: pUC19-s linearized with *Eco*RI, pUC19-s control plasmid, pUC19-s cleaved by wt BaRecJ, and pUC19-s cleaved by BaRecJ E503A mutant, H524A mutant, E560A mutant, E640A mutant, and E560A/E640A double mutant, respectively.

RecJ contain a PIWI-like domain, which is an important domain entrust chopping activity in pAgo protein. BaRecJ homologs contain similar PIWI-like regions, which have some similarities with other PIWI proteins and Ago proteins (Supplementary Figure 7). Thus, we tested the activities of homologs of BaRecJ, including BhRecJ, BoRecJ, FeRecJ, and BcRecJ. When we induced expression of these proteins, we found that they also led to death of the host *E. coli* (Supplementary Figure 8). These results imply that these homologs of BaRecJ are, like BaRecJ itself, endonucleases. It can be inferred that the C-terminal domain which contains PIWI-like domain give rise to the DNA chopping activity. Some of the PIWI domains of Agos have an RNase H-like fold, and their active sites contain a DEDX (X = N, D or H) catalytic tetrad, which confers endonuclease activity on these proteins (25, 26). We predicted the 3D structure of BaRecJ through the RCSB PDB website (http://www.rcsb.org/#Subcategory-search sequences). It showed that the residues of the DEDX motif was corresponding to D561, E600, L634 and S762 in BaRecJ (Supplementary Figure 9). *In vitro*, the D561A mutant disrupted the ability of the enzyme to digest plasmid DNA, while the L634A mutant increased the activity, and the mutants of the other two positions had little effect on the activity (Figure 5d, e).

The increase in activity caused by the mutation may be because it is more favorable for binding to the substrate or catalytic reaction, or the steric hindrance is reduced after the mutation of the inactive site, which is beneficial to the binding of the substrate. In addition, we mutated other positions based on the predicted structure and sequence comparisons. The E640A mutant showed reduced cleavage activity, however, other three mutants E503A, H524A, E560A did not affect the activity (Figure 5f). These results imply that residues D561 and E640 are important for the endonuclease activity of BaRecJ.

### BaRecJ can cleave plasmid by using small DNA guides

Our data show that BaRecJ has endonuclease activity, so how does it cut? We used different guides (25 nucleotides [nt]) for DNA-guided cleavage reactions (Supplementary Figure 10A-C). General and phosphorylated 25-nt single guides resulted in degradation of pUC-19s, as was also observed in the absence of any guide. Use of an S-modified guide resulted in no degradation of pUC19-s plasmid, but a reduction in the amount of supercoiled plasmid and an increase in linearization were obvious. Considering that the protein also has exonuclease activity and the results obtained in DNA-guided cleavage reactions using DNA guides completely matching the target plasmid (Supplementary Figure 10A-C), we decided to test S-modified DNA guides of different lengths and for different reaction times. It was found that with increasing S-modified guide length (15, 20, 25 and 35 nt), the rate and extent of degradation of the plasmid were reduced (Supplementary Figure 10D, E). As the reaction time was prolonged, the proportion of open circular and linear plasmid DNA increased (Supplementary Figure 10A-C). Additionally, L634A, the activity increased mutant, also showed improved ability to produce open circular and linear plasmid DNA after reloaded S-modified DNA guides (Supplementary Figure 11). These results indicating that the 25-nt S-modified DNA guide could inhibit the chopping activity of BaRecJ, but enhance the ability of BaRecJ to linearize the plasmid.

To test the specificity of the cleavage position, we used a 25-nt S-modified ssDNA guide with BaRecJ to treat pUC-19s for 1 and 4 h, respectively, and recovered linearized fragments. After ligation, transformation and sequencing, it was found that the mutation (i.e. deletion/insertion) sites were near the guide in the short-term reaction (Figure 6a, b), indicating that the 25-nt S-modified ssDNA guide can bind to the active site and help cleave the plasmid. When the reaction time was prolonged, the mutation site was further away from the position of the guide, and the length of the deletion fragment was increased (Figure 6c, d). It can be concluded that general guides are degraded by BaRecJ, while S-modification decreases degradation of the guide by BaRecJ.

**Figure 6.**
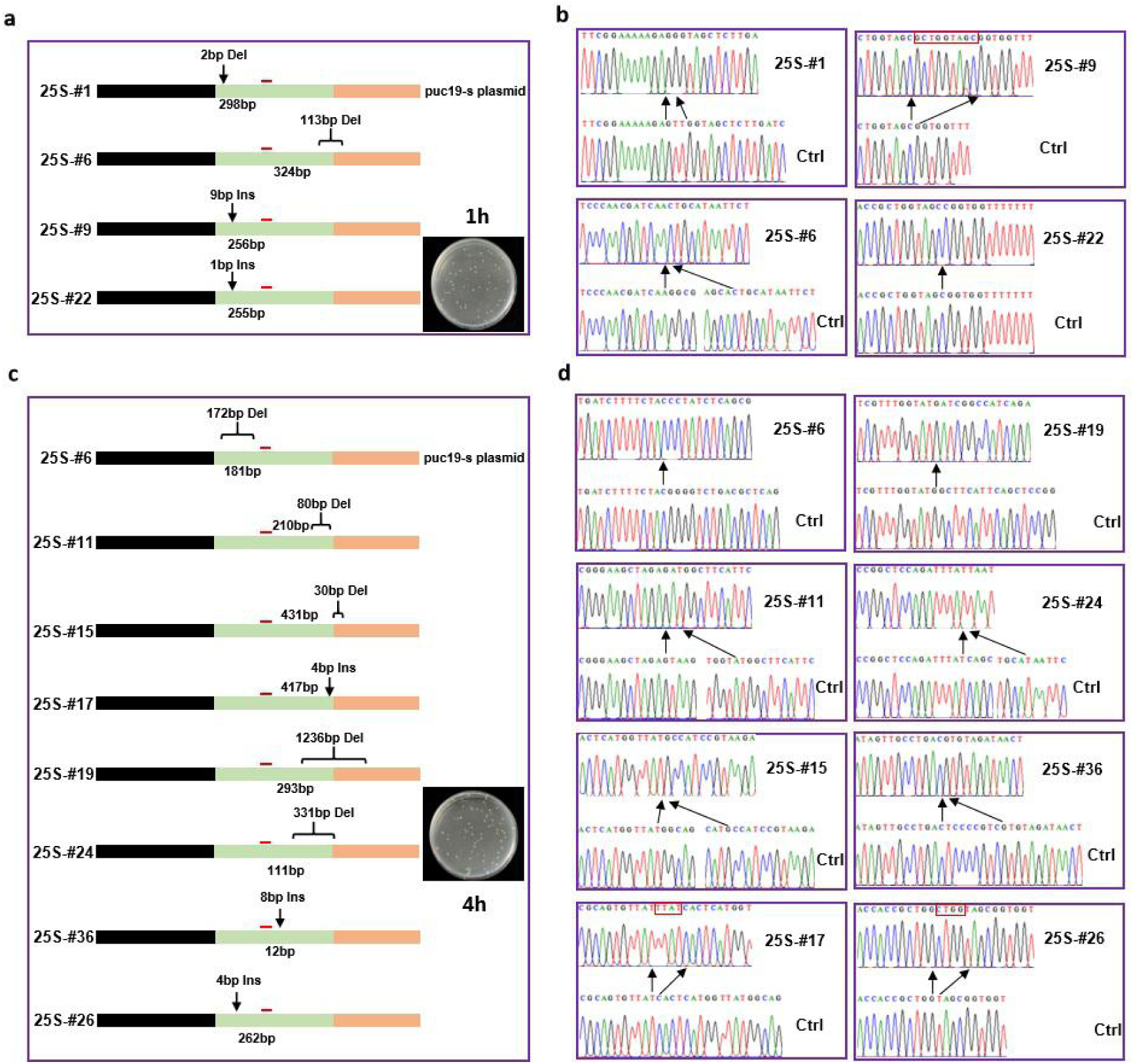
Sequence change of plasmid cleaved by BaRecJ using a guide. **a** Examples of sequencing results after DNA-guided cleavage of pUC19-s DNA, which shows that deletions and insertions were randomly produced near the guide/target region (500 nM BaRecJ, 10 μg plasmid DNA, cleavage for 1 h at 37°C). **b** Representative sequencing results after 1 h of cleavage. **c** Examples of sequencing results after DNA-guided cleavage of pUC19-s DNA, which show that deletions and insertions were randomly produced near the guide/target region (500 nM BaRecJ, 10 μg plasmid DNA, cleavage for 4 h at 37°C). **d** Representative sequencing results after 4 h of cleavage. The above results are from 60 independent experiments. Represents the guide site.

*In vivo*, BaRecJ can directly interact with plasmid or genomic DNA and cause DNA DSBs, leading to a decrease of DNA content. *In vitro*, BaRecJ can cleave and degrade circular plasmid DNA. Without a guide DNA, BaRecJ can directly cause ‘chopping’ (27) and degradation of a plasmid. BaRecJ may be able to degrade these DNAs in a non-specific fashion via guide-independent endonuclease function, similar to that of several Ago proteins, such as the archaeal MjAgo (28); besides canonical guide-dependent endonuclease activity, MjAgo has guide-independent endonuclease activity. This kind of ‘chopping’ activity would allow BaRecJ to cleave long dsDNA, including circular plasmid DNA and genomic DNA. The resulting cleavage products may then be used as guides for further cleavage of the starting substrate.

Due to the exonuclease activity of BaRecJ, when no modified guide is bound to BaRecJ, the guide can be digested from the terminus, which causes the target plasmid to be cleaved in a guide-independent manner. Thus, it was difficult to verify guide-dependent specific endonuclease function using a general guide *in vitro*. We thus observed the characteristics of the endonuclease activity by undertaking experiments using S-modified guides instead; such modification can inhibit digestion of the guide to some extent. When the guide length reached 25-nt, its digestion by BaRecJ was significantly inhibited, improving the ability of the enzyme to cleave a specific target sequence. Therefore, such modified short DNA fragments can be used as a guide to direct BaRecJ to cleave a specific site. The DHH and DHHA1 domains of BaRecJ bind to the 5ʹ- end of a ssDNA. Based on the principle of base complementation, we suggest that BaRecJ binds to the target strand in a position complementary to the single-stranded guide DNA. BaRecJ then nicks the target single strand of the plasmid, which is bound by the guide, and further nicks the other strand to linearize the plasmid. The linearized fragment is further degraded in the direction from 5ʹ- to 3ʹ-, to finally form a sticky end protruding at the 3ʹ- end.

BaRecJ has a different structure from typical Ago and PIWI proteins (27) and other types of RecJ protein (15). Compered with the typical EcRecJ, BaRecJ also possesses a DHH domain and a DHHA1 domain. In addition, BaRecJ possesses a novel PIWI-like domain at the C-terminus, which is different from other Ago and PIWI proteins (27, 28). It does not have a conserved DEDX tetrad with endonuclease activity (25,26,29); only the first two residues are conserved—the latter two residues are different from those in the other proteins. BaRecJ exerts endonuclease activity, but its guide-dependent cleavage mechanism is fundamentally different from Agos. It lacks the MID and PAZ domains, which usually form binding pockets that facilitate the anchoring of the 5ʹ- and 3ʹ-ends of an oligonucleotide guide, respectively. In contrast, BaRecJ anchors the guide through the DHH domain and is then directed to the specific cleavage site.

In *B. alcalophilus*, canonical RecJ domains and a PIWI-like domain are coupled together to function, but in other bacteria these two proteins are separate. However, it is possible that they work together through protein interactions. Whether BaRecJ is exceptional, or whether genes encoding RecJ and PIWI-like proteins were originally fused but separated and then evolved separately, deserves further study. RecJ and PIWI-like proteins may participate in repair or defense processes through interactions. In particular, the eukaryotic RecJ homologue, Cdc45, which only has ssDNA binding capability (13), is worthy of further study. We believe that BaRecJ contains the classical DHH and DHHA1 domains, which can exert the exonuclease function and cut off mismatched bases, and play a role in the process of DNA mismatch repair. At the same time, BaRecJ, which binds to the mismatch repair site, can also function as an endonuclease to cleave exogenous plasmid DNA or viral DNA near the mismatch repair site, to protect the repair site from interference from foreign DNA and ensure the smooth progress of repair.

As a unique branch of the DHH superfamily, RecJ nucleases with a PIWI-like domain at the C-terminus are widely present in class Bacilli. In future, more research is required for complete understanding of the functional diversity and differentiation of RecJs that have an additional C-terminal domain. These C-terminal domains generally show sequence similarity to Ago proteins, such as the PIWI-like domain in BaRecJ. Here, our results support a novel function of the C-terminal domain in restricting exogenous plasmids. BaRecJ is the first reported RecJ with endonuclease and exonuclease activity, and is possibly involved in maintaining genome stability. BaRecJ can perform DNA-guided cleavage of target DNA *in vitro* at a wide range of temperatures; it functions efficiently at 37°C. Therefore, it can potentially be used for genome editing and selective cleavage of dsDNA targets from cells of mesophilic organisms. A putative mechanism of action by BaRecJ was proposed (Figure 7).

**Figure 7.**
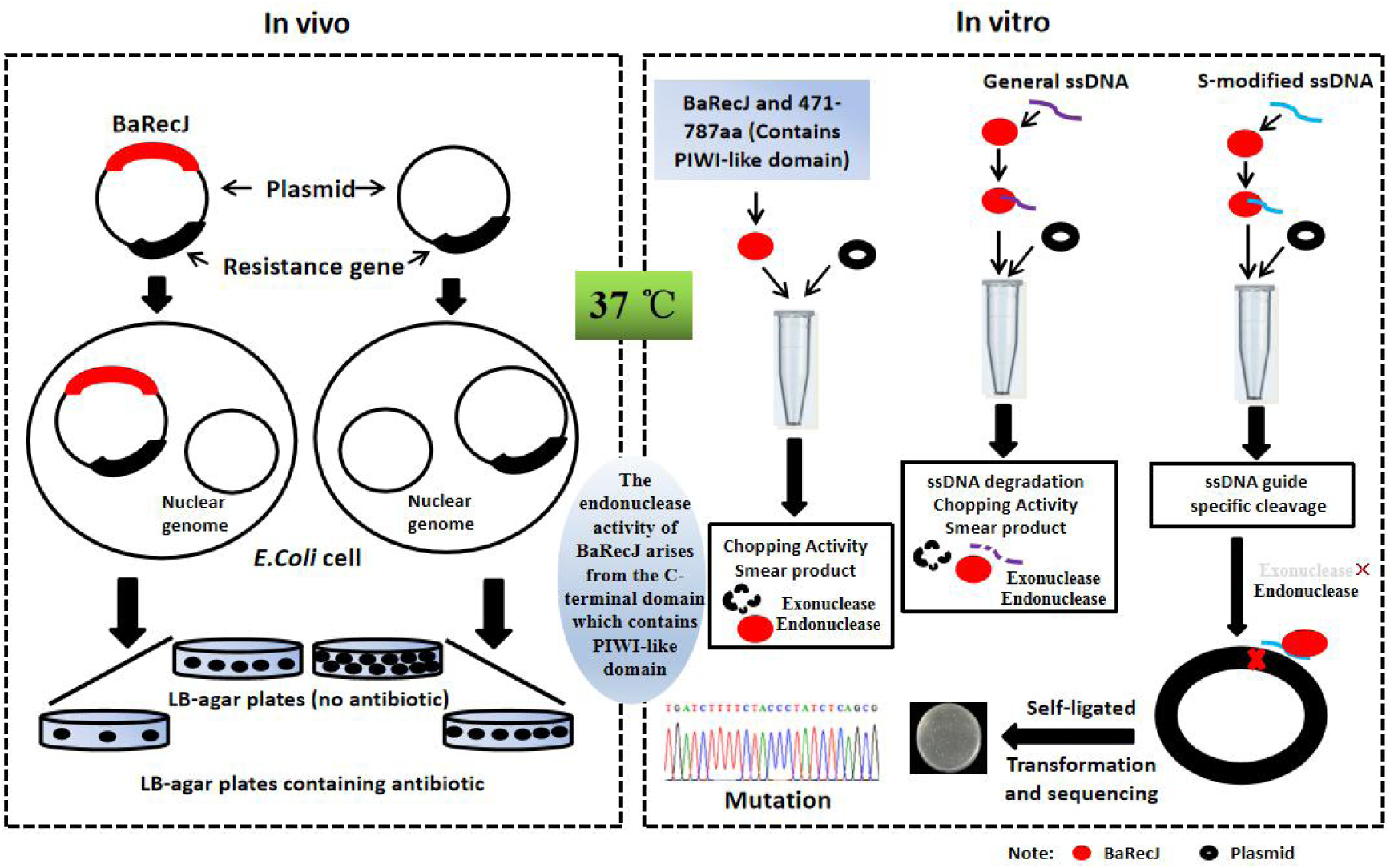
A putative mechanism of action by BaRecJ

## CONCLUSION

In this study, we find that BaRecJ is extremely different from other DHH superfamily members [25, 26]. It uniquely exhibits endonuclease activity, cleaving dsDNA and causing mutation and even cell death. According to the experiment in vitro, BaRecJ cleaves supercoiled plasmid into open circular and linear plasmid, and completely degrades the supercoiled plasmid into nucleotides. The activity of BaRecJ is influenced by temperature and the concentration of Mg2+. While the influence of other metal ions remains to be further studied. In vivo the basal level of BaRecJ (without induction) makes cells form the long filaments but not influences the cell survival; in contrast, a high content of BaRecJ (after IPTG induction) influences the cell survival, causing the cell death and less filaments. Moreover, BaRecJ cleaves specific target sequences at moderate temperature when directed by a phosphorothioate-modified single-stranded DNA (S-modified ssDNA) guide. These findings suggest that BaRecJ is substantially different from any reported RecJs and we believe that once the regulation mechanism of BaRecJ is deciphered, it can be applied in diverse fields in the future.

## Experimental Procedures

### Culture of *Bacillus alcalophilus*

*B. alcalophilus* CGMCC 1.3604 (ATCC27647) was purchased from the China General Microbiological Culture Collection Center and cultivated in maltose/peptone/yeast extract medium (1.0% maltose, 0.5% peptone, 0.1% yeast extract, 0.1% K_2_HPO_4_, 0.02% MgSO_4_·7H_2_O, 1.5% Na_2_CO_3_) at 30°C and pH 9.7.

### Plasmid construction and transformation

The gene encoding BaRecJ (aa 1–787) was amplified by PCR from *B. alcalophilus* genomic DNA (Supplementary Table 1). PCR product was cloned into pMD18-T and confirmed by DNA sequencing. The *BarecJ* DNA fragment was further subcloned into constitutive expression vector pET-23b(+) between *Nde*I and *Xho*I sites, yielding pET23BaRecJ, and into inducible vectors pACYCDuet-1 and pET28a between *Nde*I and *Xho*I sites, generating pACYCBaRecJ and pET28BaRecJ respectively. Plasmids for expressing site-directed mutants of BaRecJ were constructed using phosphorothioate-based ligase-independent gene cloning, as previously described (30). RecJs from other species (Bh, Bc, Bo, Fe, Sg) were treated in the same way.

### Analysis of transformation efficiency

*E. coli* BL21 (DE3) competent cells were spread on LB-agar plates containing 100 μg ml^−1^ ampicillin and 50 μg ml^−1^ chloramphenicol to select cells harboring pET-23b(+) and pACYCDuet-1 respectively. Transformation efficiency of pET-23b and pET23BaRecJ was calculated on the basis of the number of colony-forming units (CFU) per µg DNA. To compare transformation efficiency, the number of CFU for pET23BaRecJ was defined as 1, with the number of CFU for pET-23b(+) normalized to this value. The sequence of plasmids from surviving transformants was confirmed by traditional Sanger sequencing by Personal Biotechnology Company (Shanghai, China). Primers used are listed in Supplementary Table 2.

### DNA content assay

*E. coli* BL21 (DE3) containing pACYCDuet-1 or pACYCBaRecJ were cultivated in LB medium supplemented with 50 μg ml^−1^ chloramphenicol. The culture was collected after induction for 3 h and used to extract plasmid and genomic DNA using an E.Z.N.A.® Plasmid Mini Kit I (Omega) and an E.Z.N.A.® Bacterial DNA Kit (Omega), respectively, according to the user manuals. After preliminary confirmation of the quality of the DNA on a 0.8% agarose gel, the plasmid and genomic DNAs were quantified using a BioTeke ND5000 spectrophotometer. RecJs from other species (Bh, Bc, Bo, Fe) were assessed in the same way.

### Bacterial survival assay

*E. coli* BL21 (DE3) colonies containing pACYCBaRecJ or empty vector (pACYCDuet-1) were selected to prepare suspensions, respectively. Appropriate volumes of suspensions were spread onto LB agar plates containing 0.1 mM IPTG and 50 μg ml^−1^ chloramphenicol, and cultivated at 37°C; then CFU were counted. To identify the effect of BaRecJ on survival rate, the number of CFU for plasmid pACYCBaRecJ was defined as 1, and the number of CFU for pACYCBDuet1-1 was normalized based on the value for pACYCBaRecJ. The effect of induction time on survival rate was studied by inducing bacteria harboring pACYCDuet-1 or pACYCBaRecJ with IPTG for increasing periods of time. The colonies were inoculated into LB medium containing 0.1 mM IPTG and 50 μg ml^−1^ chloramphenicol and cultivated at 37°C with shaking. Samples were taken every hour for 8 h. The samples were centrifuged at 13,000 × *g* for 3 min and washed three times at room temperature to remove IPTG. The resuspensions were spread on LB-agar plates, cultivated at 37°C overnight, then CFU per ml were counted. Transformants containing pET28BaRecJ and its mutants or empty vector (pET28a) were selected and grown to the same OD_600_. The diluted bacterial suspensions of different strains were spotted onto LB-agar containing 50 μg ml^−1^ kanamycin and 0.1 mM IPTG and cultured overnight at 37°C.

### Expression and purification of BaRecJ and its mutants

Recombinant expression plasmids were transformed into *E. coli* BL21 (DE3) competent cells (Supplementary Table 3). Transformants were cultivated in LB medium containing 50 µg ml^−1^ kanamycin and 50 μg ml^−1^ chloramphenicol at 37°C in a shaking incubator and grown to an OD_600_ of 0.5–0.6. The cells were cold-shocked by incubating in an ice bath for 20 min. Protein expression was induced for 16 h at 16°C by adding IPTG to a final concentration of 0.2 mM. Cells were harvested by centrifugation. Recombinant proteins were purified via immobilized Ni^2+^-affinity chromatography as follows: the bacterial pellet was suspended in lysis buffer (50 mM Tris-HCl, pH 8.0, 0.5 M NaCl, 20 mM imidazole) and disrupted by sonication. The cell extract was clarified by centrifugation at 13,000 × *g* for 30 min. The supernatant was loaded onto a resin-containing column pre-equilibrated with lysis buffer and washed with >100 column volumes of lysis buffer containing 20 mM imidazole. The bound protein was eluted from the column using elution buffer (50 mM Tris-HCl, pH 8.0, 0.5 M NaCl, 250 mM imidazole). The affinity-purified proteins were polished by gel-filtration chromatography using Superdex G200. After checking the purity of the eluate by 15% SDS-PAGE, the protein was dialyzed into storage buffer (50 mM Tris-HCl, pH 8.0, 0.15 M NaCl and 30% glycerol) and stored in small aliquots at −80°C.

### Nuclease activity assays

Nuclease activity toward dsDNA was characterized as described with some modifications^15, 26^. Plasmid DNA, PCR-amplified linear DNA, or genomic DNA was used as substrate. Except in the titration assay of BaRecJ, all reactions were performed with a concentration ratio of substrate to BaRecJ of 1:20 at 37°C. Titration of plasmid to BaRecJ was performed with concentration ratios 1:1, 1:5, 1:10, 1:20, and 1:40. To determine the metal ion preference of BaRecJ, digestion of plasmid DNA by BaRecJ was performed at different concentrations of Mg^2+^ (0.1, 1, 5, 10, 50 mM) in buffer (10 mM Tris-HCl, pH 8.0, 50 mM NaCl and 10 mM dithiothreito (DTT)) at 37°C for 2 h. After optimization of the divalent ion concentration, the reaction buffer consisted of 10 mM Tris-HCl, pH 8.0, 50 mM NaCl, 10 mM MgCl_2_ and 10 mM DTT. The reactions were stopped by adding an equal volume of 50 mM ethylenediaminetetraacetic acid (EDTA), and incubated for 15 min at 65°C. Reaction products were analysed on 1% agarose gels. Gels were stained with 4S Red Plus Nucleic Acid Stain (BBI Life Sciences), visualized under a GelDoc XR system, and analysed using Image Lab software (Bio-Rad).

Nuclease activity toward single-stranded oligonucleotides and oligoribonucleotides, as well as their double-stranded forms (Supplementary Table 4), was characterized similarly to the method described above for dsDNA. The reactions were performed at 37°C for 30 min in reaction buffer consisting of 10 mM Tris-HCl, pH 8.0, 50 mM NaCl, 10 mM MgCl_2_ and 10 mM DTT. After incubation, an equal volume of stopping buffer (90% formamide, 100 mM EDTA and 0.2% sodium dodecyl sulfate) was added to the reactions. Then, the reaction mixtures were subjected to 8 M urea-denaturing 15% PAGE to separate the products. Finally, the gels were imaged and quantified using an FL9500 fluorescence scanner (GE Healthcare). EMSA experiments were used to characterize the binding capability of BaRecJ and the BaRecJ H170A mutant to oligonucleotides and oligoribonucleotides, as well as their double-stranded forms. During EMSA assays, MnCl_2_ in the buffer was replaced with CaCl_2_, and 10% native PAGE was run to detect potential protein–substrate complexes.

Supercoiled plasmid DNA was used as the target in DNA-guided cleavage assays (Supplementary Table 5). The reactions were performed at molar ratio 1:20:30 target:protein:guide at 37°C for 1 or 4 h. BaRecJ (500 nM) was mixed with 750 nM guide DNA in reaction buffer consisting of 10 mM Tris-HCl, pH 8.0, 50 mM NaCl, 10 mM MgCl_2_ and 10 mM DTT, and incubated at 37°C for 20 min for guide reloading. The target plasmid DNA was added to a final concentration of 25 nM. Reactions were analysed on 1% agarose gels. Gels were stained with 4S Red Plus Nucleic Acid Stain, visualized under a GelDoc XR system and analysed using Image Lab software. The obtained linearized plasmid was excised from the agarose gel and purified using a MiniBEST Agarose Gel DNA Extraction Kit (Takara Bio). The cohesive ends of the linearized plasmid were blunt-ended and self-ligated using a DNA Blunting Kit (Code No. 6025, Takara Bio). The DNA Blunting Kit can smooth DNA with 3ʹ- or 5ʹ- protruding ends; T4 DNA polymerase, one of components in the kit, has 5ʹ- to 3ʹ- polymerase activity and 3ʹ- to 5ʹ- exosomal enzyme activity. The resulting blunt-ended DNA can be efficiently ligated using the ligation solution in the kit. The ligation product was then transformed into competent *E. coli* DH5α cells (Takara Bio), and after 16 h of culture, selected clones were sent for sequencing (Sangon Biotech).

### Statistical analysis

All P-values were calculated by Student’s *t*-test using SPSS software version 22.0. Statistical significance was accepted at the 95% confidence level, which meant that a difference was considered significant if P < 0.05. The P-values of transformation efficiencies were calculated by inputting the number of CFU from biological triplicates of plasmid-bearing cells. In survival experiments, the number of CFU from biological triplicates of transformants cultivated on the plate was used as the input. The P-values for plasmid and genome DNA content were obtained by inputting the plasmid and genome DNA yields from biological triplicates of transformants after 3-h induction.

## ACKNOWLEDGMENTS

This work was supported by the National Key R&D Program of China (Grant No. 2018YFC0310701), the Natural Science Foundation of China (Grant No. 31671539, 31370214), the Stem Cell Strategy Library and Clinical Transformation Platform of Stem Cell Technology in Shanghai Zhangjiang National Independent Innovation Demonstration Zone (ZJ2108-ZD-004).

---

**Supplementary Figure 1.**
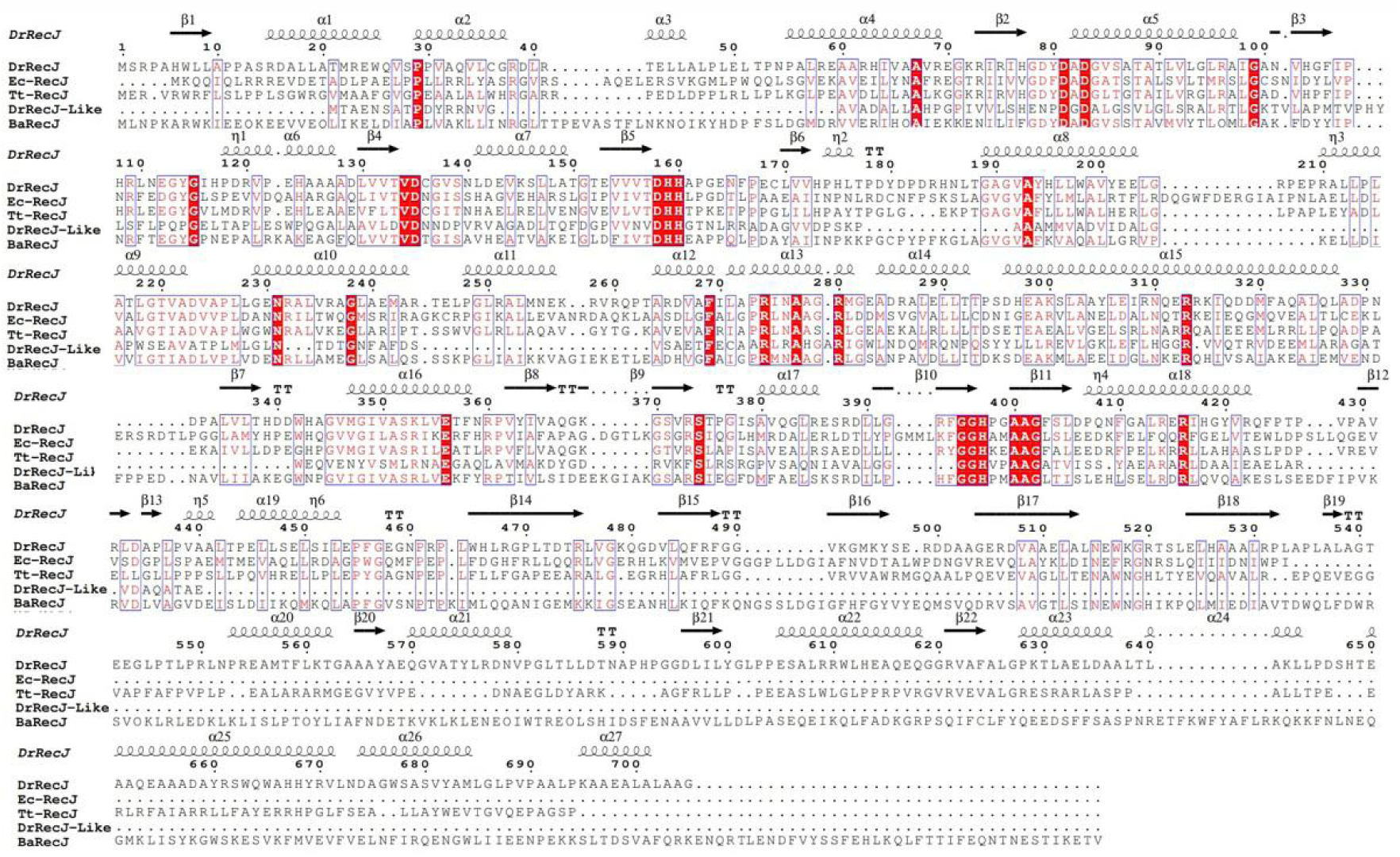
Multiple sequence alignment of bacterial RecJs and sequence analysis of mutations in BaRecJ. Sequence alignment of several representative RecJ/DHH protein superfamily members. Ba, Dr, Ec and Tt are *Bacillus. alcalophilus*, *Deinococcus. radiodurans*, *Escherichia. coli, Thermus. thermophilus*, respectively. Conserved domains, DHHA1 and DHH, are highlighted with frames.

**Supplementary Figure 2.**
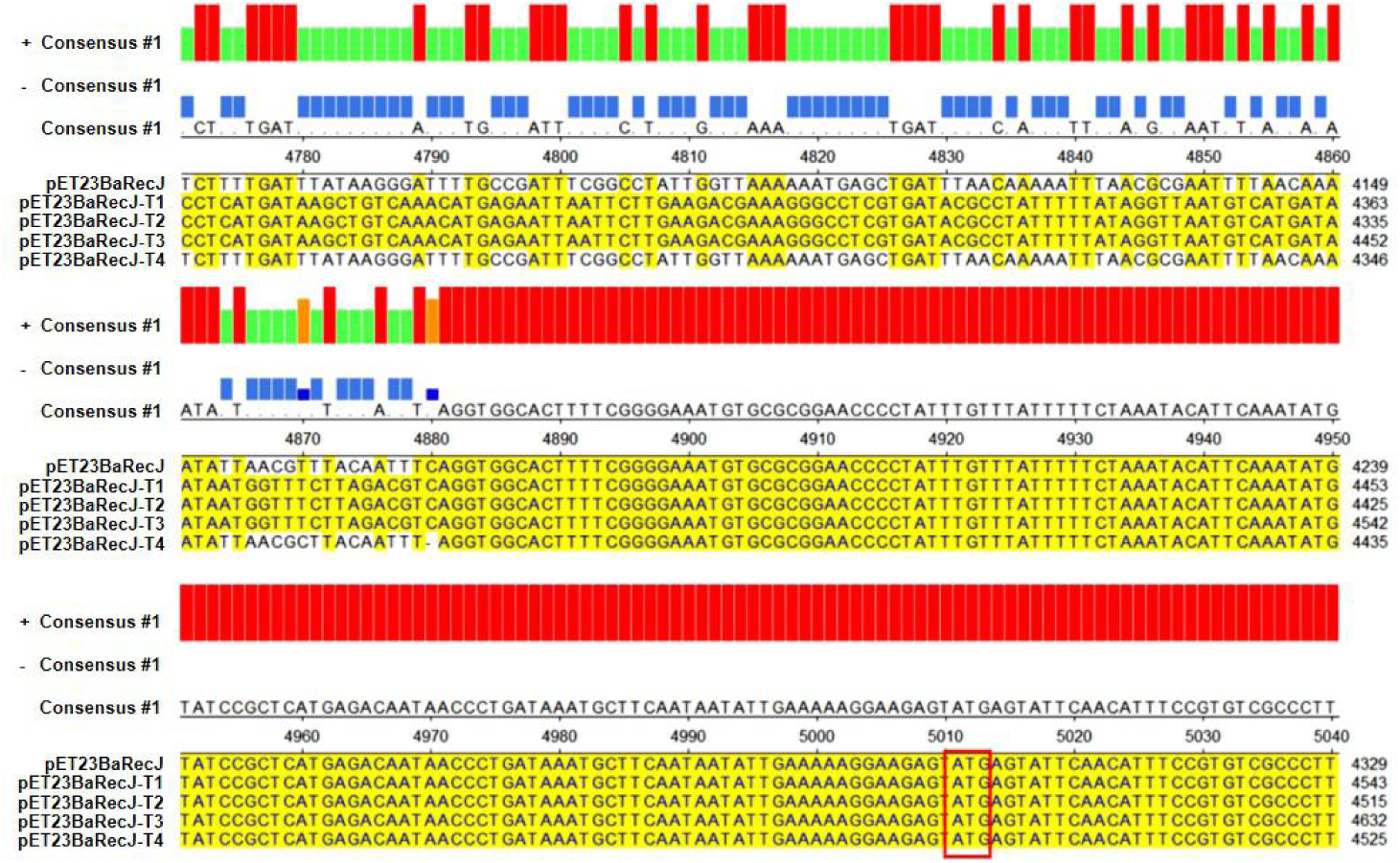
Comparison of sequences between pET23BaRecJ and pET23BaRecJT1 to T4. The plasmids pET23BaRJT1 to T4 were respectively isolated from surviving transformants of pET23BaRJ and sequenced. The DNA region from bases 4000 to 5100 was aligned. The ATG start codon of the antibiotic resistance gene is highlighted in a black rectangle.

**Supplementary Figure 3.**
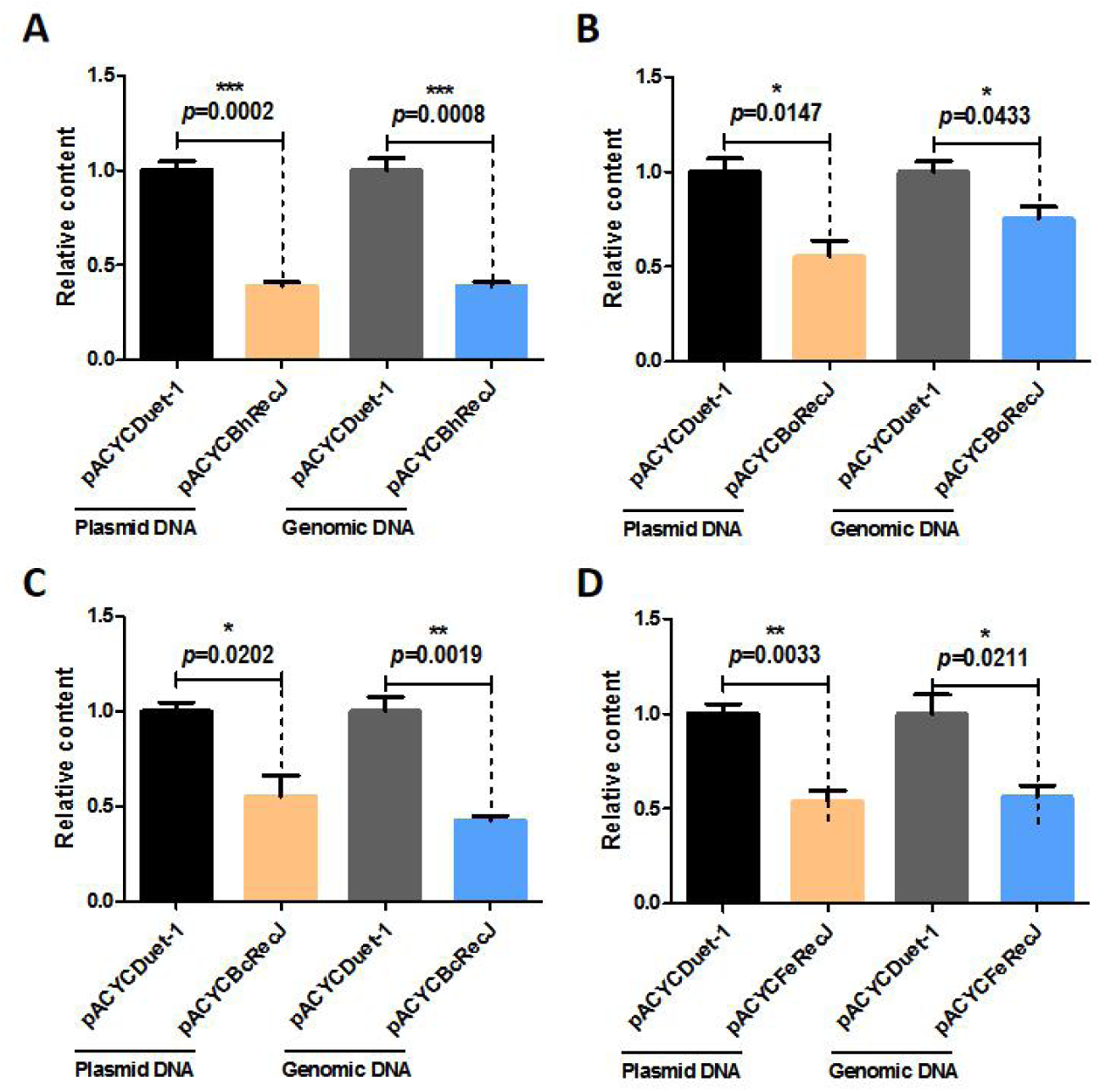
Plasmid and genome content of *E. coli* BL21 (DE3) transformed with BaRecJ homologs. **A** Yield of plasmid and genomic DNA from cells transformed with pACYCBhRecJ or pACYCDuet-1. **B** Yield of plasmid and genomic DNA from cells transformed with pACYCBoRecJ or pACYCDuet-1. **C** Yield of plasmid and genomic DNA from cells transformed with pACYCBcRecJ or pACYCDuet-1. **D** Yield of plasmid and genomic DNA from cells transformed with pACYCFeRecJ or pACYCDuet-1. The plasmid and genomic DNAs were extracted from the same amount of biomass of transformants induced for 3 h. All data were obtained from mean values of three independent experiments. Error bars depict standard deviations of biological triplicates.

**Supplementary Figure 4.**
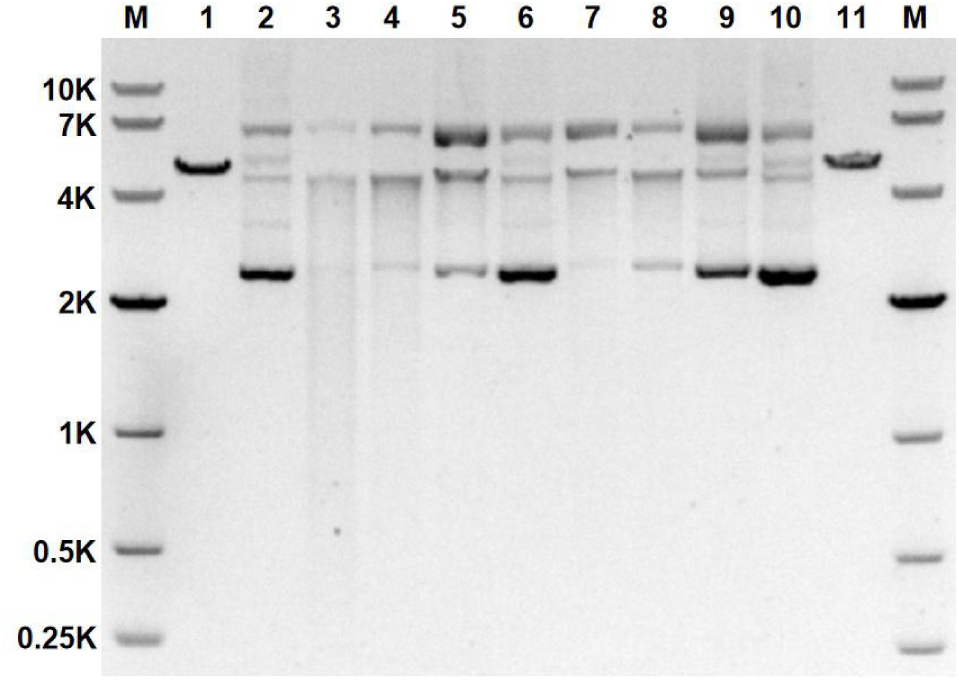
The metal preference of BaRecJ. The activity of BaRecJ (500 nM BaRecJ, 750 ng plasmid DNA, cleavage for 2 h at 37°C) in the presence of 1 mM Mg^2+^, Mn^2+^, Ca^2+^, Cu^2+^, Li^+^, Ni^2+^, or Co^2+^ (lanes 3–9, respectively). Lane 1, pUC19-s control linearized with *Eco*RI; lane 2, pUC19-s plasmid control.

**Supplementary Figure 5.**
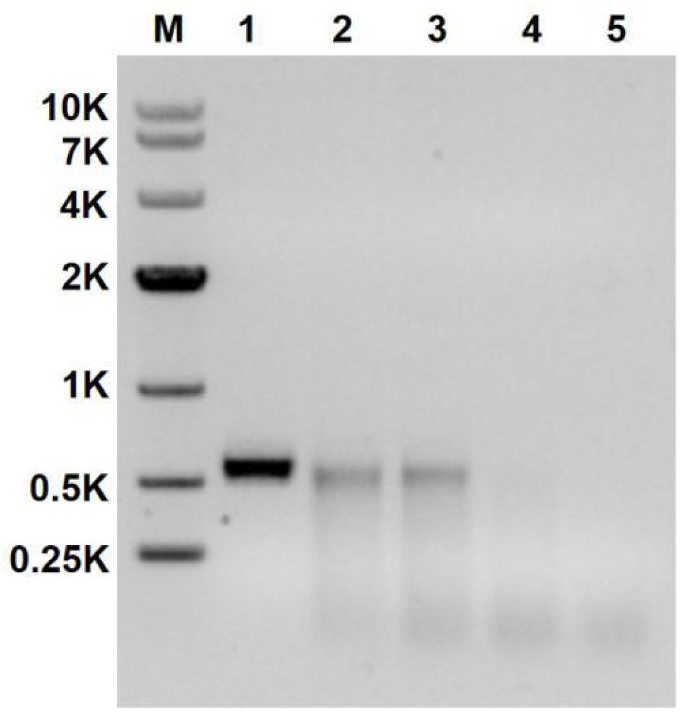
Time course of BaRecJ activity toward linear dsDNA (593 bp) at 37°C (500 nM BaRecJ, 750 ng linear dsDNA). Lanes 1–5: linear dsDNA control, and digestion for 30, 60, 120, and 180 min, respectively.

**Supplementary Figure 6.**
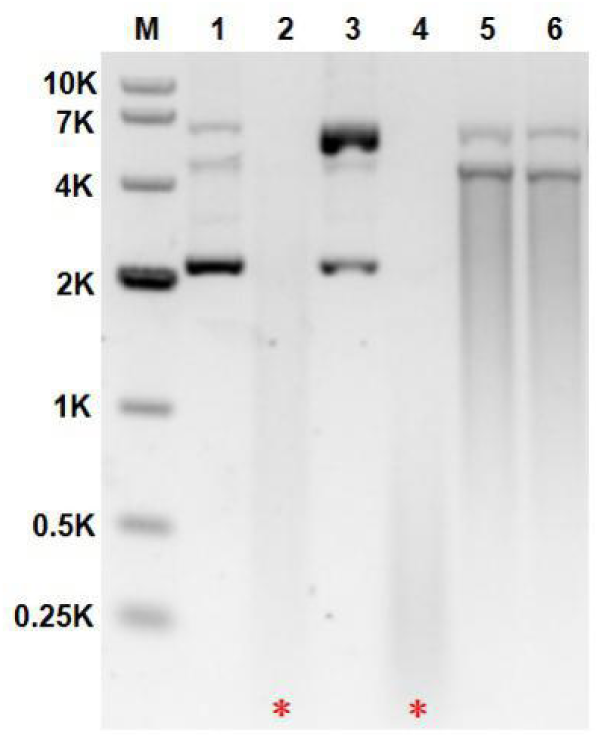
Electrophoresis of the final plasmid degradation products of wt BaRecJ and its truncations (500 nM protein, 750 ng plasmid DNA, cleavage for 2 h at 37°C). Lanes 1–6: pUC19-s control plasmid, cleavage by wt BaRecJ, and cleavage by BaRecJ fragments aa 1–471, aa 471–787, aa 471–653, and aa 653–787, respectively.

**Supplementary Figure 7.**
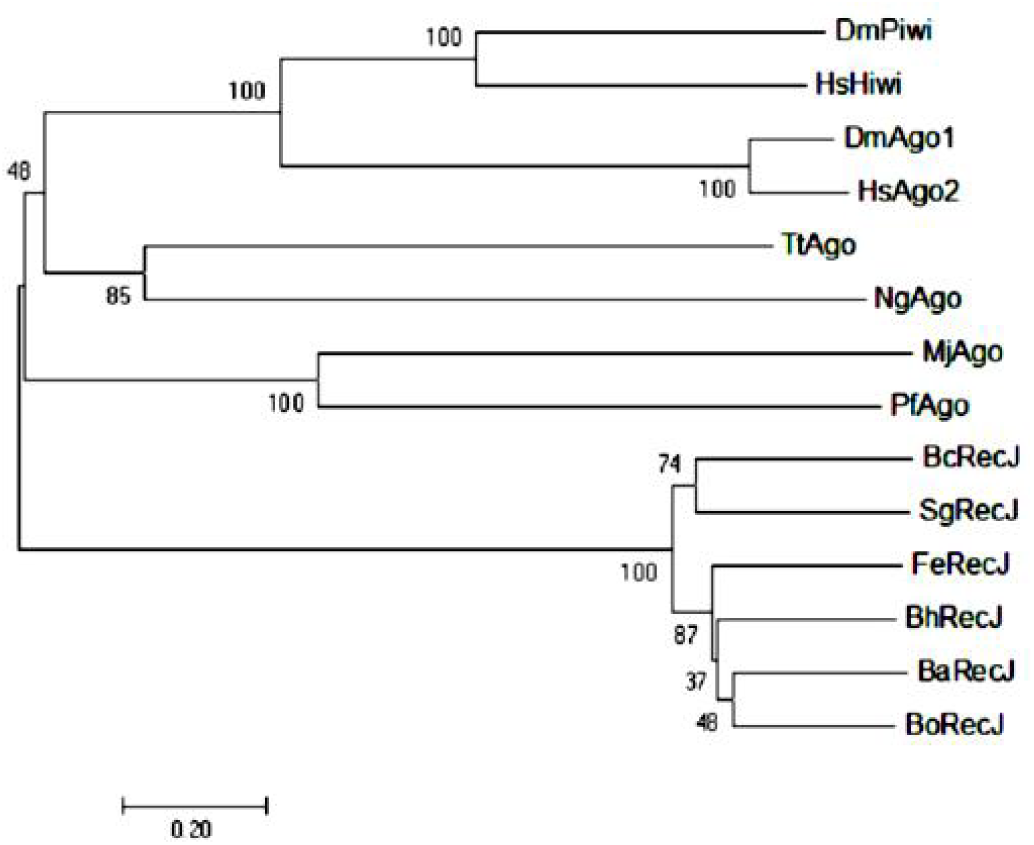
Phylogenetic tree of bacterial RecJs. The bootstrap values are listed at the branch points.

**Supplementary Figure 8.**
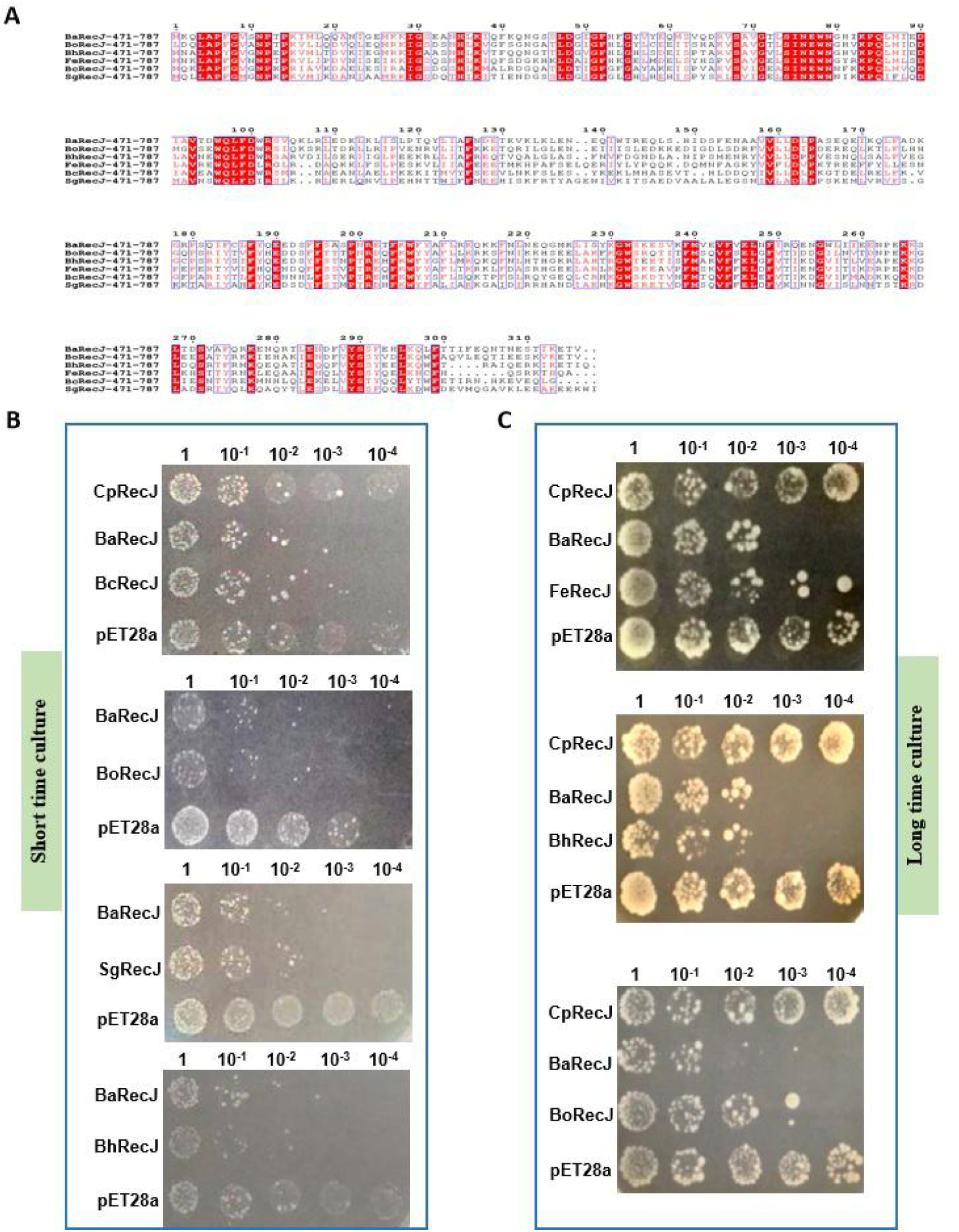
Comparison of RecJ with homologous proteins. **A** Sequence alignment of several representative RecJ protein superfamily members. Ba, Bo, Bh, Fe, Bc, Sg are *B. alcalophilus*, *B. okhensis*, *B. halodurans*, *F. enclensis*, *B. cereus*, and *Sporosarcina globispora*, respectively. **B, C** The activity of wt BaRecJ and its homologous proteins *in vivo*. Grown to the same OD_600_, diluted bacterial suspensions of wt BaRecJ, cells transformed with homologous genes, and a pET28a(+) vector control strain were spotted onto plates containing 0.1 mM IPTG and cultured overnight at 37°C.

**Supplementary Figure 9.**
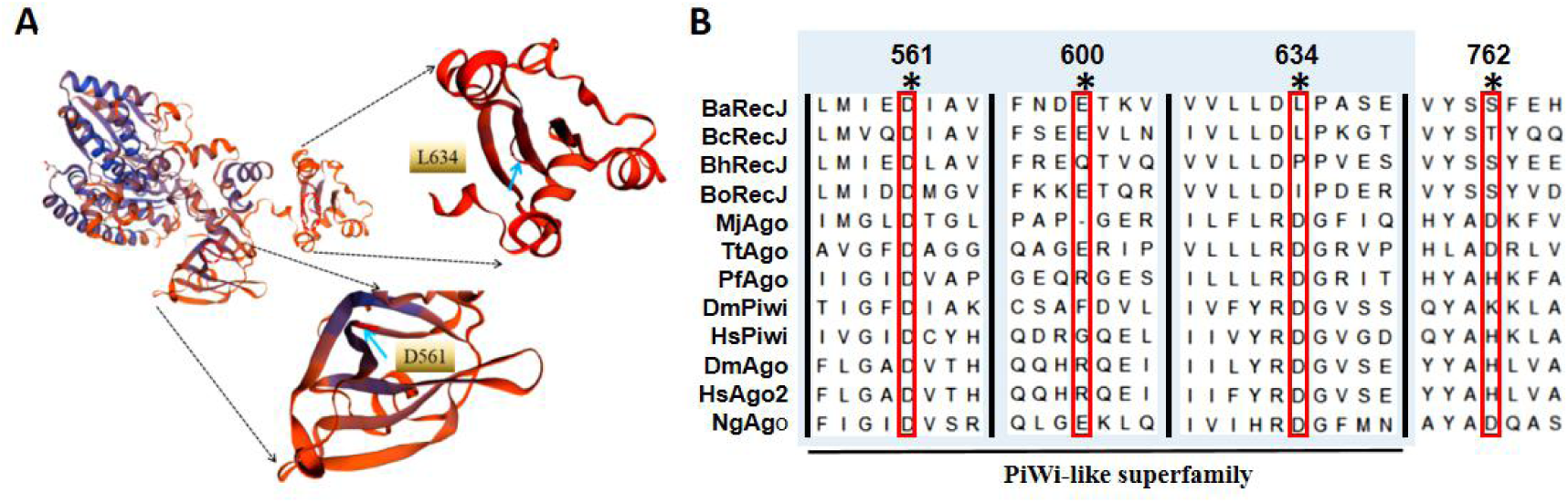
Bioinformatics analysis of BaRecJ. **A** Prediction of 3D structure of BaRecJ through RCSB PDB website. **B** Multiple sequence alignment including the core motifs of BaRecJ and homologous proteins. Red boxes denote predicted active-site residues of wt BaRecJ.

**Supplementary Figure 10.**
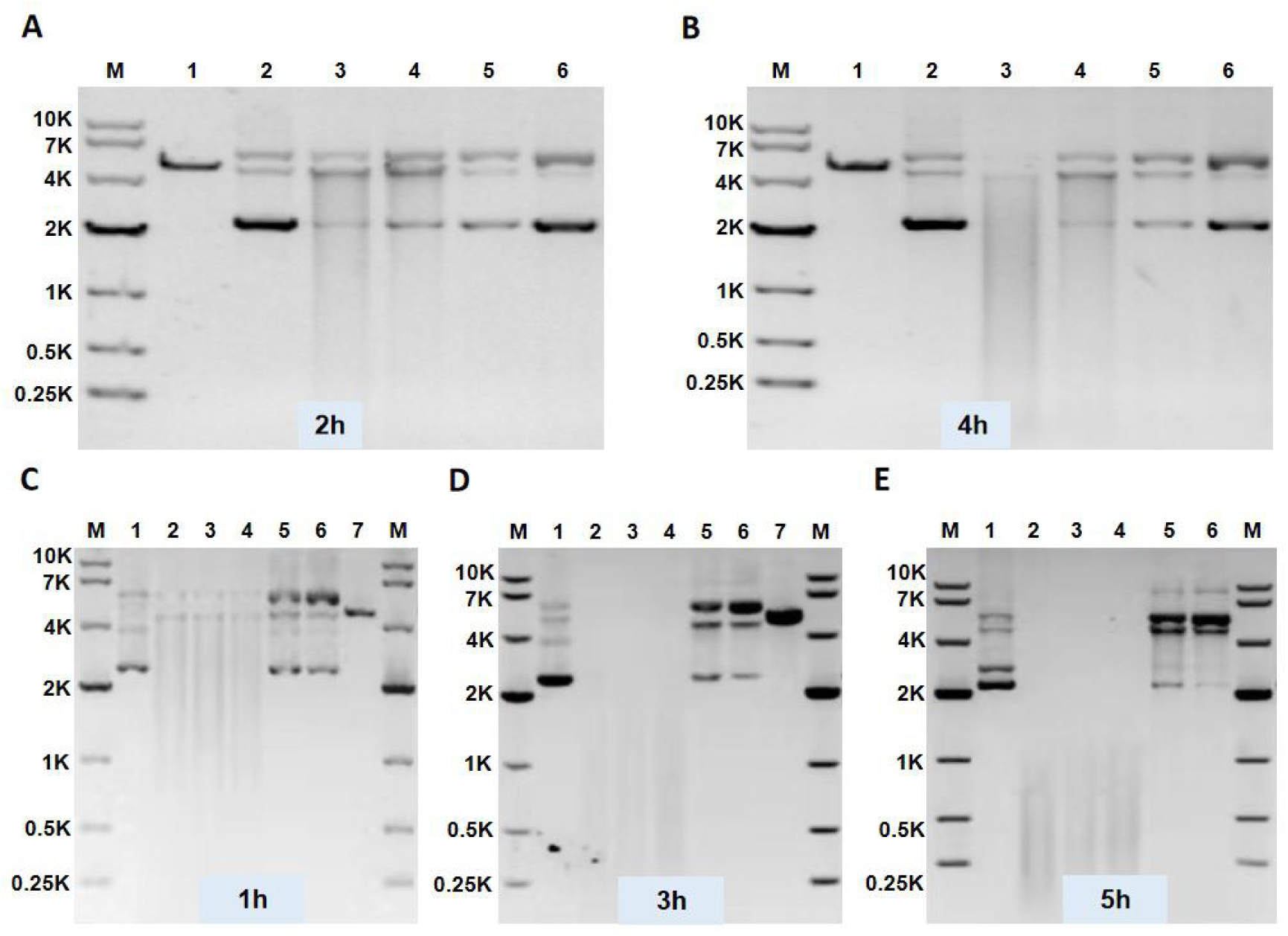
Characterization of the nuclease activity of BaRecJ reloaded with different guides. **A**, **B** and **C** Different guides (25 nucleotides [nt]) were used for DNA-guided cleavage reactions; 800 nM BaRecJ was reloaded with 1.2 μM ssDNA guide, mixed with 40 nM target plasmid DNA and incubated at 37°C for 1 (**A**), 3 (**B**), or 5 h (**C**). Lanes 1–7: pUC19-s control plasmid, cleavage by wt BaRecJ, effect of BaRecJ reloaded with 5ʹ-P ssDNA, effect of BaRecJ reloaded with 5ʹ-OH ssDNA, effect of BaRecJ reloaded with S-modified ssDNA, effect of BaRecJ reloaded with S-modified 5ʹ-P ssDNA, and pUC19-s linearized with *Eco*RI. **D, E** Different guide strand lengths (15, 20, 25, 35 nucleotides [nt]) were used in cleavage reactions (800 nM BaRecJ, 1.2 μM S-modified ssDNA guide, 40 nM target plasmid DNA). The reactions were incubated at 37°C for 2 (**D**) or 4 h (**E**). Lanes 1–6: pUC19-s linearized with *Eco*RI, pUC19-s control plasmid, pUC19-s cleaved by wt BaRecJ and 15-nt S-modified ssDNA guide, pUC19-s cleaved by wt BaRecJ and 20-nt S-modified ssDNA guide, pUC19-s cleaved by wt BaRecJ and 25-nt S-modified ssDNA guide, pUC19-s cleaved by wt BaRecJ and 35-nt S-modified ssDNA guide, respectively.

**Supplementary Figure 11.**
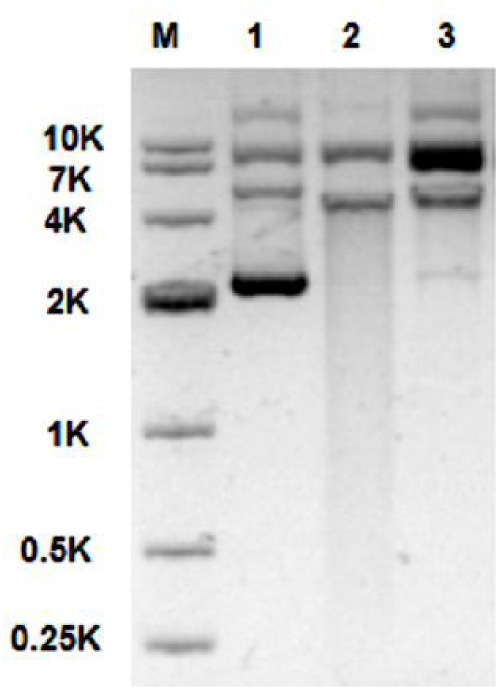
Characterization of the nuclease activity of BaRecJ L634A mutant reloaded with S-modified ssDNA guide. 500 nM BaRecJ L634A mutant was reloaded with 1 μM ssDNA guide, mixed with 25 nM target plasmid DNA and incubated at 37°C for 15min. Lanes 1–3: pUC19-s control plasmid, pUC19-s cleaved by BaRecJ L634A mutant, pUC19-s cleaved by reloaded BaRecJ L634A mutant.

**Supplementary Table 1.**
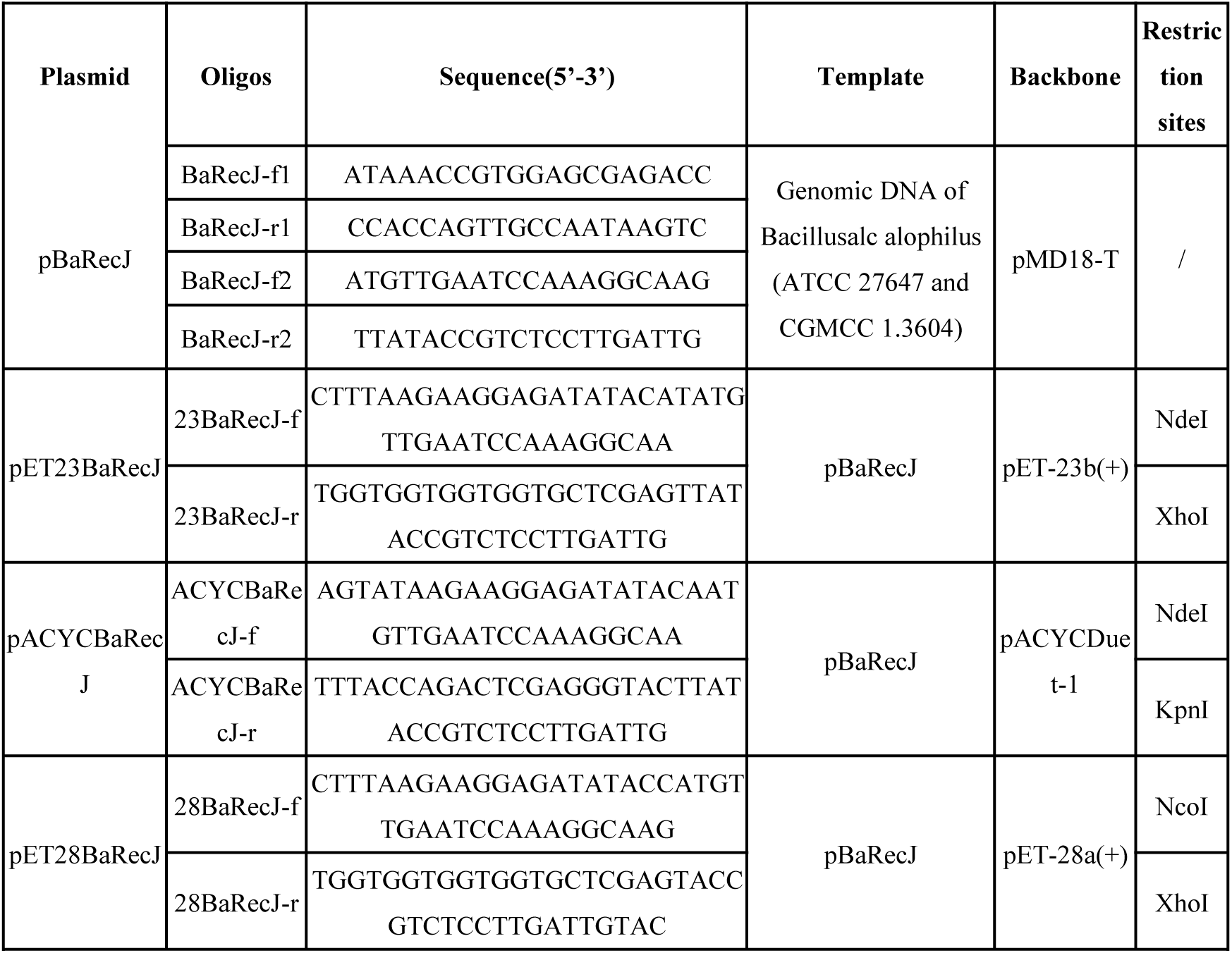
Primers, plasmids used in this experiment.

**Supplementary Table 2.**
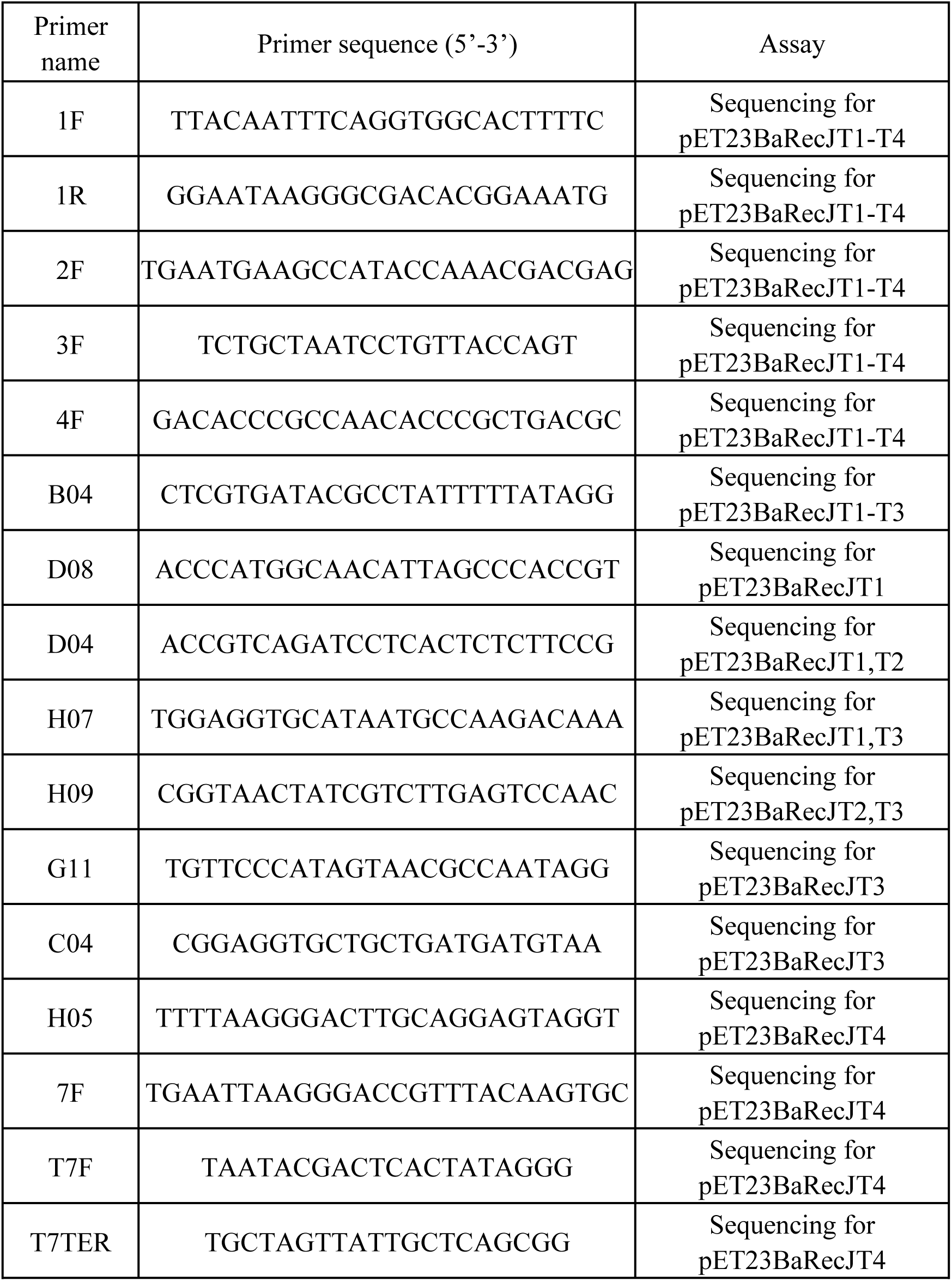
Primers used for plasmid sequencing.

**Supplementary Table 3.**
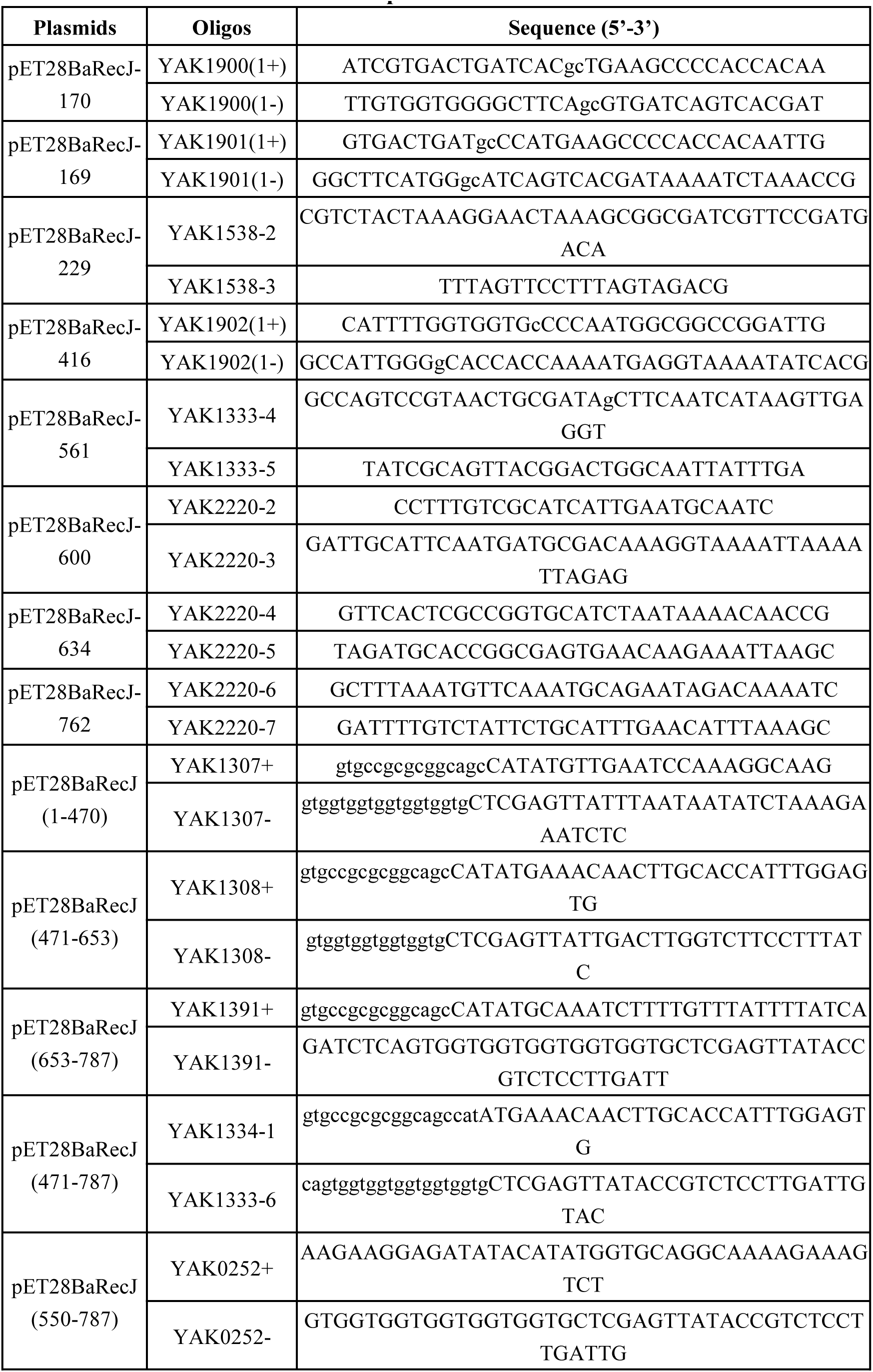
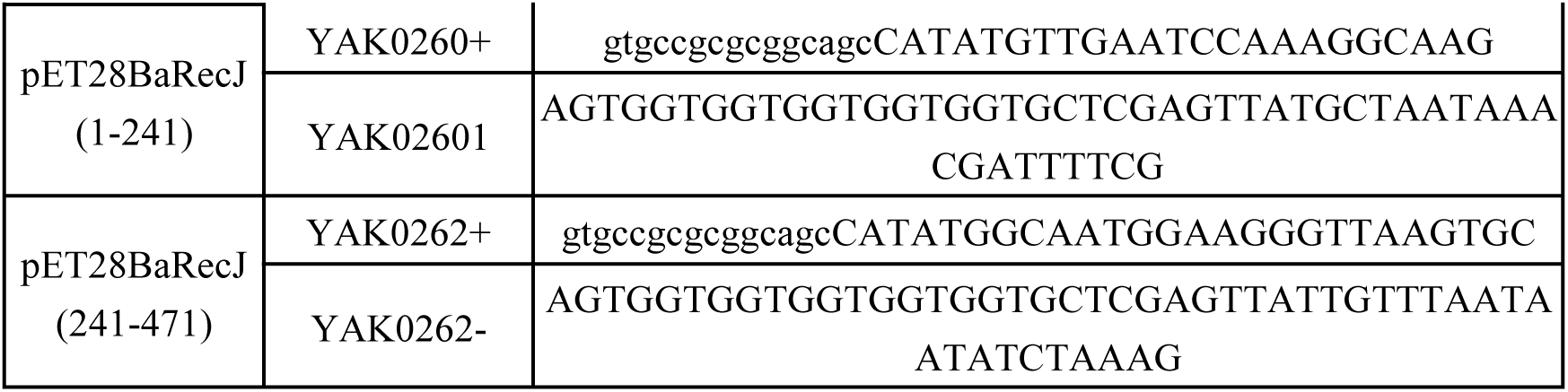
Primers used for constructing BaRecJ mutation plasmids.

**Supplementary Table 4.**
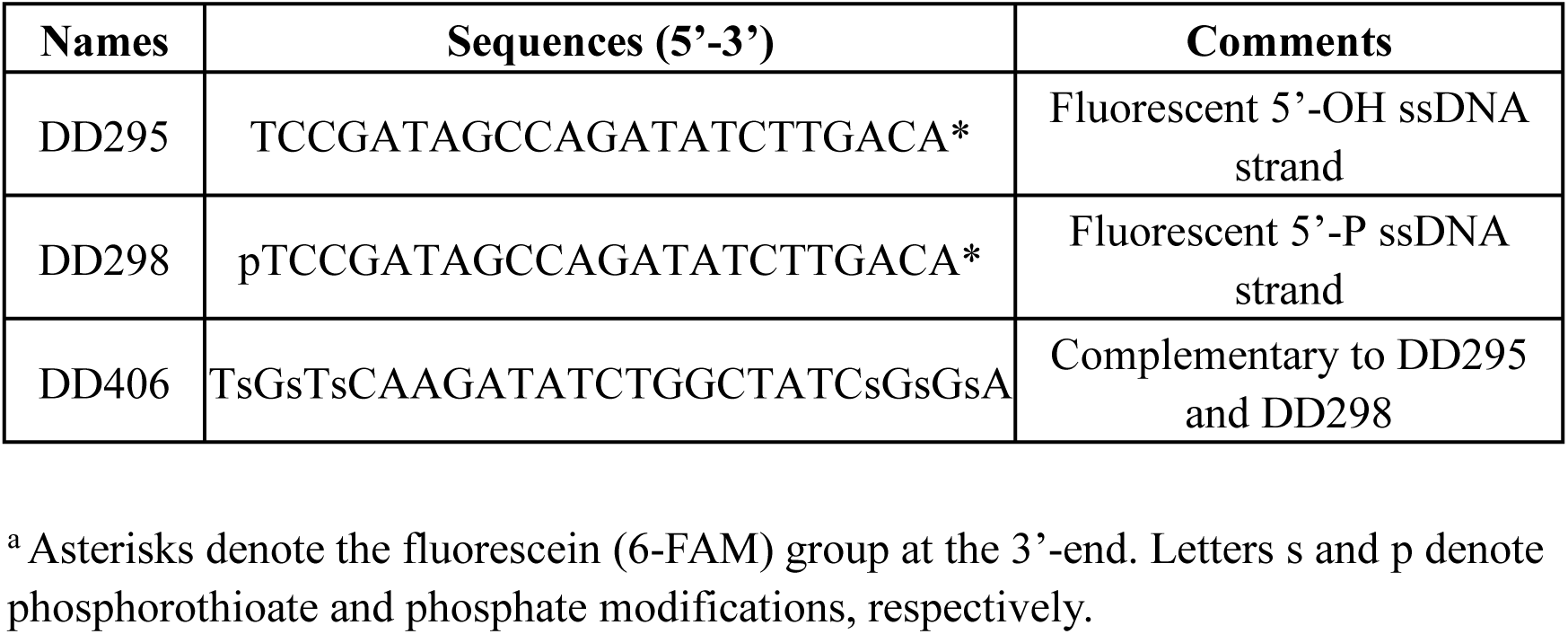
Oligonucleotides used for characterizing the nuclease activity.

**Supplementary Table 5.**
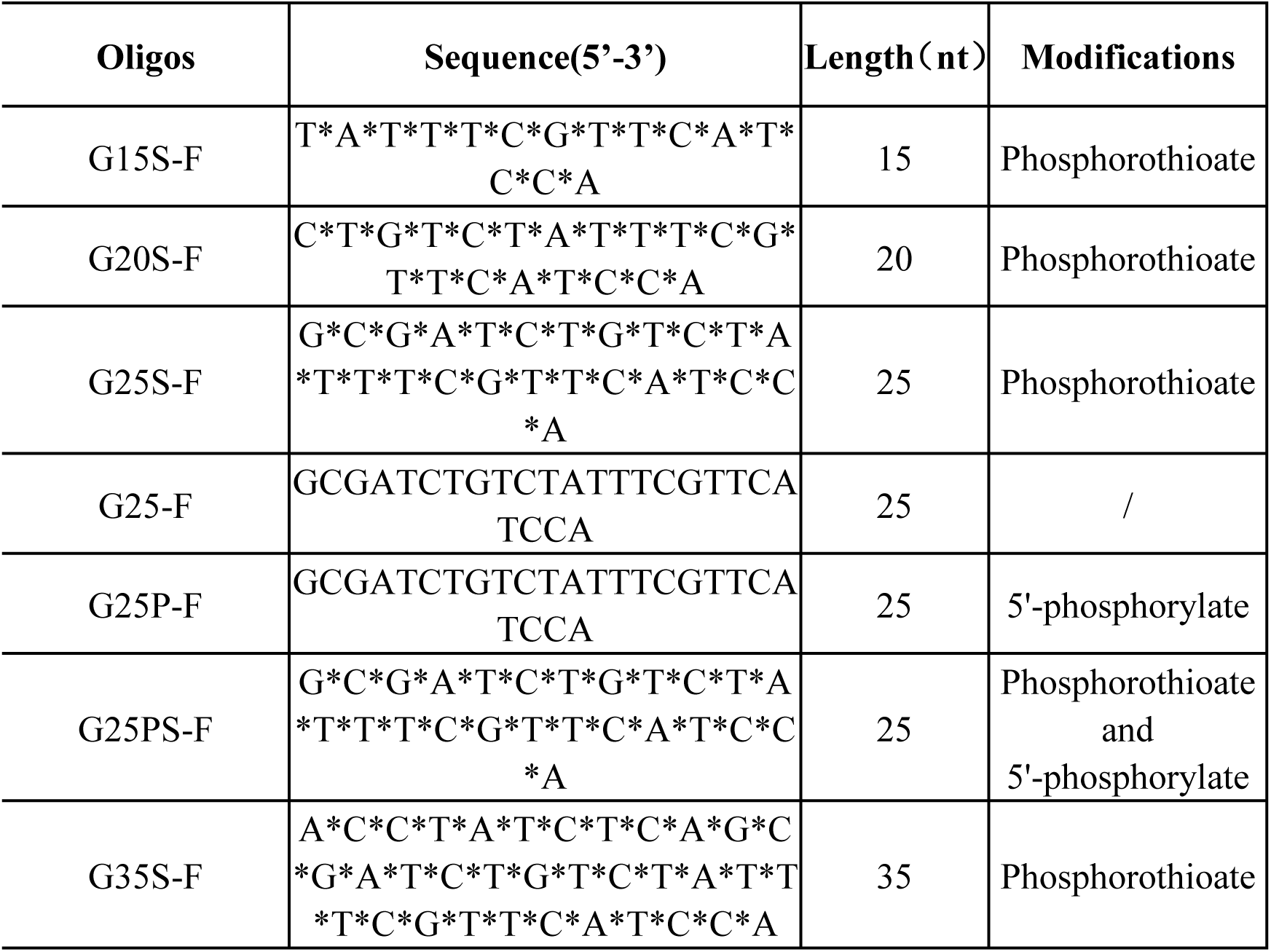
Oligos used for characterizing endonuclease activity of BaRecJ.

